# Nanopore sequencing for real-time genomic surveillance of *Plasmodium falciparum*

**DOI:** 10.1101/2022.12.20.521122

**Authors:** Sophia T. Girgis, Edem Adika, Felix E. Nenyewodey, Dodzi K. Senoo Jnr, Joyce M. Ngoi, Kukua Bandoh, Oliver Lorenz, Guus van de Steeg, Alexandria J. R. Harrott, Sebastian Nsoh, Kim Judge, Richard D. Pearson, Jacob Almagro-Garcia, Samirah Saiid, Solomon Atampah, Enock K. Amoako, Collins M. Morang’a, Victor Asoala, Elrmion S. Adjei, William Burden, William Roberts-Sengier, Eleanor Drury, Megan L. Pierce, Sónia Gonçalves, Gordon A. Awandare, Dominic P. Kwiatkowski, Lucas N. Amenga-Etego, William L. Hamilton

## Abstract

Malaria is a global public health priority causing over 600,000 deaths annually, mostly young children living in Sub-Saharan Africa. Molecular surveillance can provide key information for malaria control, such as the prevalence and distribution of antimalarial drug resistance. However, genome sequencing capacity in endemic countries can be limited. Here, we have implemented an end-to-end workflow for *P. falciparum* genomic surveillance in Ghana using Oxford Nanopore Technologies, targeting antimalarial resistance markers and the leading vaccine antigen *circumsporozoite protein* (*csp*). The workflow was rapid, robust, accurate, affordable and straightforward to implement, and could be deployed using readily collected dried blood spot samples. We found that *P. falciparum* parasites in Ghana had become largely susceptible to chloroquine, with persistent sulfadoxine-pyrimethamine (SP) resistance, and no evidence of artemisinin resistance. Multiple Single Nucleotide Polymorphism (SNP) differences from the vaccine *csp* sequence were identified, though their significance is uncertain. This study demonstrates the potential utility and feasibility of malaria genomic surveillance in endemic settings using Nanopore sequencing.

## Introduction

Malaria is a major cause of morbidity and mortality worldwide, particularly for young children living in Sub-Saharan Africa. The World Health Organization (WHO) estimates that there were 247 million malaria cases and 619,000 deaths in 2021 [1]. 76% of deaths were among children under 5 years old and 95% were in Africa [1]. The COVID-19 pandemic disrupted essential malaria control services, setting back the progress made in the 2000-2019 period [1]. The WHO has identified antimalarial drug resistance as a key threat to control and elimination efforts [2]. Artemisinin-based Combination Therapy (ACT) is the current front-line treatment for *Plasmodium falciparum* malaria – the most virulent species responsible for the majority of deaths. ACT is highly effective and well tolerated, and has been a cornerstone of recent progress in reducing the burden of malaria disease worldwide. Artemisinin partial resistance has been defined as delayed clearance of parasites carrying specific mutations following treatment with an artemisinin derivative despite adequate dosing and absorption [2]. In combination with partner drug resistance, artemisinin partial resistance can cause treatment failure [3]. If such parasites become widespread in Africa, the results would be devastating.

The capacity for parasite populations to undergo evolutionary change requires ongoing surveillance to monitor for new threats. Having first emerged and spread in Southeast Asia [3–12], artemisinin partial resistance – caused by mutations in the gene *kelch13* [13–15] – has now been identified in Rwanda [16–18], Uganda [19,20] and Eritrea [2], and appears to have emerged independently in Africa and Southeast Asia [16]. Resistance to Sulfadoxine-pyrimethamine (SP), caused by mutations in the target genes *dhfr* and *dhps* [21–25], threatens the efficacy of intermittent preventive therapy in pregnancy (SP-IPTp) and seasonal malaria chemoprevention (SMC) in young children (used in combination with amodiaquine, SP+AQ) [26]; these are important public health interventions to protect vulnerable populations in hyperendemic regions. Parasite genome sequencing, incorporated into surveillance programmes, can provide key information to guide National Malaria Control Programme (NMCP) decision-making; for example, describing the geospatial distribution and longitudinal trends of antimalarial resistance markers [27–30] and *P. falciparum* population structure and relatedness [31–35].

An effective vaccine would be a hugely valuable addition to the malaria control armamentarium [36]. In October 2021, RTS,S/AS01 became the first malaria vaccine to be recommended by WHO for children living in areas of moderate to high *P. falciparum* transmission, and implementation is being piloted in Ghana, Malawi and Kenya, with plans to scale up over the next few years [1,37]. The RTS,S vaccine targets circumsporozoite protein (*csp*), expressed on the surface of sporozoites and required for hepatocyte invasion [38]. RTS,S/AS01 vaccine efficacy is around 36% after four doses [39]. Another *csp*-based vaccine, R21-M Matrix (MM), has been shown to provide up to 75% efficacy in an ongoing trial in Burkina Faso [40,41]. *csp* is also the target of long-acting monoclonal antibodies, which are showing promise as novel methods of protection [42–44]. It is unclear whether genetic diversity in *csp* has an impact on *csp*-based vaccines or therapeutic antibodies, and how this may change as vaccination is scaled up and parasite exposure to the vaccine increases.

The WHO ‘Strategy to respond to antimalarial drug resistance in Africa’ (November 2022) highlights the critical need for strengthened surveillance capacity, to increase technical and laboratory capacity and to expand coverage of data on antimalarial drug efficacy and resistance in Africa [2]. However, despite its huge potential for pathogen surveillance and global health more broadly [45,46], many endemic countries in Africa have limited capacity for genomic sequencing due to factors including prohibitive costs, barriers to procurement, and a lack of sequencing and computing infrastructure [47]. Oxford Nanopore Technologies (ONT) is being increasingly used for rapid sequencing, diagnostics, antimicrobial susceptibility testing and epidemiological analysis in multiple pathogens, including SARS-CoV-2 [48–50], Zika virus [51,52], Ebola virus [53], Chikungunya virus [54], *Mycobacterium tuberculosis* [55–57], and bacterial antimicrobial resistance (AMR) and clinical metagenomics [58,59,68,60–67]. ONT devices such as the MinION are portable, relatively cheap and produce sequence data in ‘real-time’, making them well-suited to resource-limited settings including in Low- and Middle-Income Countries (LMIC). The longer sequence reads generated by ONT can provide additional advantages, such as characterising highly polymorphic or repetitive sequences or complex structural rearrangements that are challenging to access with short reads [69,70].

Nanopore has previously been applied to *P. falciparum* amplicon sequencing for drug resistance genes [71,72] and to whole genome sequencing [73]. Here, we demonstrate that Nanopore can be used prospectively for real-time genomic analysis from clinical malaria samples in an endemic setting using the current latest ONT chemistry, which reports single read accuracy of ≥99% [74]. The end-to-end process was implemented in Ghana using standard molecular biology equipment, a handheld MinION and a commercially available laptop computer. A multiplexed PCR approach targeting key antimalarial drug resistance markers and almost full-length *csp* could produce actionable data rapidly, accurately and cheaply, with a turnaround time of a few days. This demonstrates the potential utility and feasibility of using Nanopore sequencing within endemic countries for targeted malaria molecular surveillance.

## Results

### Assay design and laboratory isolate validation

A multiplexed PCR was designed targeting six parasite loci, one amplicon within each of the antimalarial drug resistance-associated genes *chloroquine resistance transporter* (*crt*), *dihydrofolate reductase-thymidylate synthase* (*dhfr*), *dihydropteroate synthetase* (*dhps*), *multidrug resistance protein 1* (*mdr1*), and *kelch13*; and the vaccine target *circumsporozoite protein* (*csp*) (**Methods, Table 1**). Amplicons were readily distinguished by gel electrophoresis, allowing for a cheap and straightforward check post-PCR (**Figure 1**). A separate PCR targeted the full-length sequence of the polymorphic surface antigen *merozoite surface protein 1* (*msp1*), around 5kb in size, to further assess the potential for long Nanopore reads to access complex genomic regions. A custom informatics pipeline built in Nextflow was used for real-time analysis and variant calling, referred to as *nano-rave* (the Nanopore Rapid Analysis and Variant Explorer tool) (details in **Methods**).

**Table 1.** *P. falciparum* genes and variants targeted in amplicon assays. A multiplex PCR targeted the drug resistance marker genes (*crt*, *dhfr*, *dhps*, *mdr1* and *kelch13*) and the vaccine and monoclonal antibody target, *csp*, in a single assay. The *msp1* PCR was performed in a separate reaction. Mutations in bold with asterisk were used as key markers of antimalarial drug susceptibility phenotyping. Details on primer sequences, amplicons and antimalarial drug susceptibility inference rules are provided in **Supplementary Notes.**

| Gene name and ID in the 3D7 parasite clone | Key mutations targeted for genotyping | Associated antimalarial resistance or other phenotype |
| --- | --- | --- |
| Chloroquine resistance transporter, *crt* (PF3D7_0709000) | K76T* | Chloroquine resistance marker |
| Dihydrofolate reductase, *dhfr* (PF3D7_0417200) | N51I, C59R, **S108N\***, I164L | Pyrimethamine resistance markers |
| Dihydropteroate synthase, *dhps* (PF3D7_0810800) | S436A, **A437G\***, K540E, A581G, A613S/T | Sulfadoxine resistance markers |
| Multidrug resistance protein 1, *mdr1* (PF3D7_0523000) | N86Y, N86F, Y184F | No direct inferences, but associated with resistance to several antimalarials including lumefantrine |
| *kelch13* (PF3D7_1343700) | Different mutations in the propeller domain, e.g. C580Y | Artemisinin partial resistance markers |
| Circumsporozoite protein, *csp* (PF3D7_0304600) | SNPs in the C-terminal Region (CTR); assess full-length consensus sequence | Leading vaccine and monoclonal antibody target antigen |
| Merozoite surface protein 1, *msp1* (PF3D7_0930300) | Assess full-length consensus sequence | Previously explored as a vaccine candidate; potential for use as a marker of complexity of infection |

**Figure 1.**
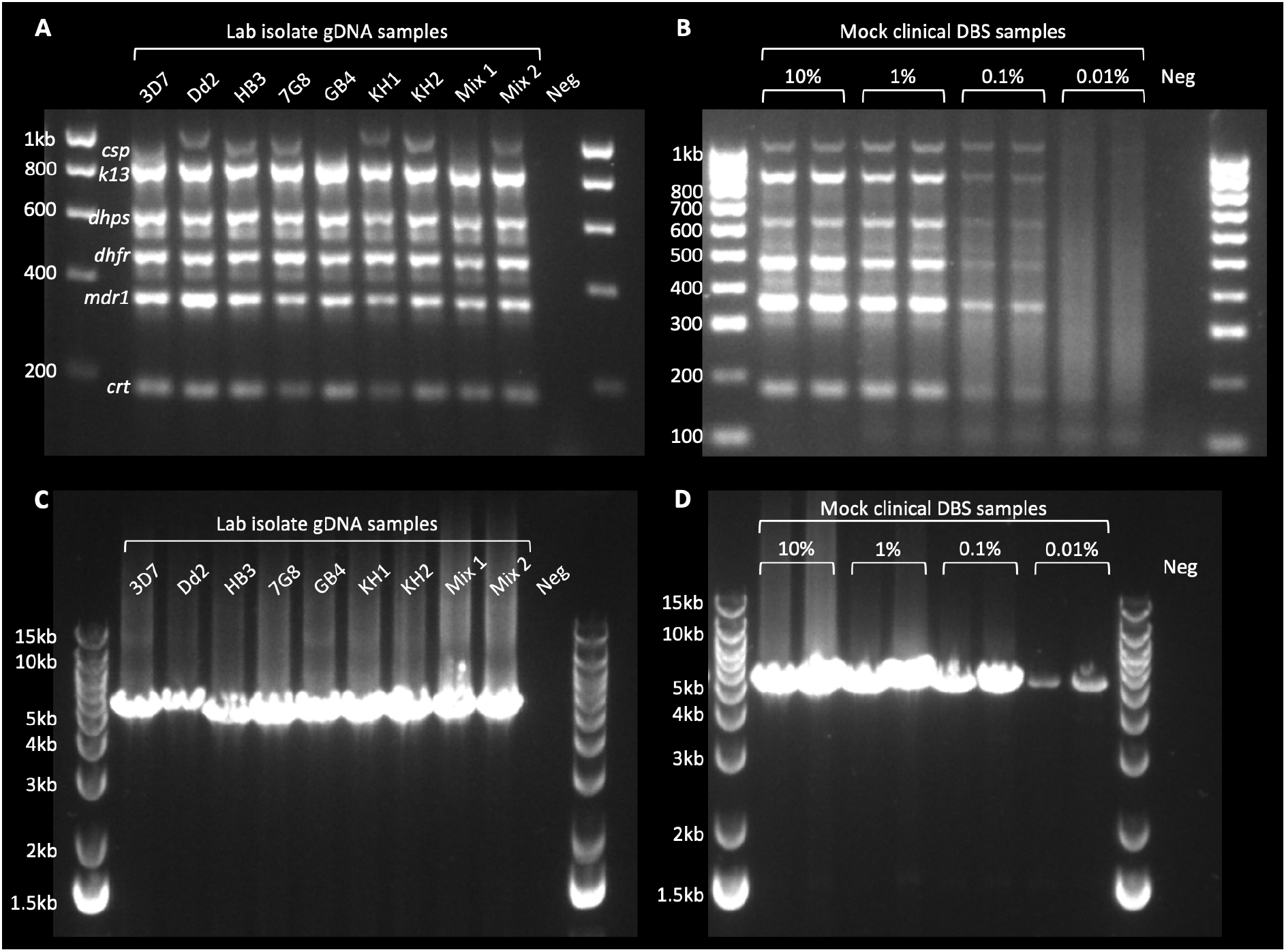
Gel electrophoresis of PCR products from laboratory clones and mock clinical samples. **A)** Multiplex drug resistance and *csp* PCR, for a selection of laboratory clones, run on 2% agarose gel. Bands are annotated based on expected sizes for each amplicon. Note variable size of *csp* due to a deletion in the N-Terminal Domain in 3D7 and variation in the Central Repeat region. Mixtures 1 and 2 contained, respectively: 3D7 + KH2 (80:20) and KH2 + 3D7 (80:20). **B)** Multiplex drug resistance and *csp* PCR, for mock clinical DBS samples, run on 2% agarose gel. Mock clinical DBS were prepared by combining *in vitro* cultured *P. falciparum* RBCs with human whole blood, in ratios expected to produce final parasitaemias of 10%, 1%, 0.1% and 0.01% infected RBCs, with 50μl blotted onto filter papers to mimic clinical DBS. The proportions of human and parasite DNA per sample were assessed by quantitative PCR (**Supplementary Figure 1**). Samples were extracted and assessed in duplicate. Although bands stopped being visible in the 0.01% parasitaemia samples on this gel, Nanopore sequence coverage was still adequate for drug resistance genotyping. **C)** *msp1* PCR, for a selection of laboratory clones, run on 1% agarose gel. A single fragment of approximately 5Kb is expected. The same samples were used as template DNA as in gel (A). **D)** *msp1* PCR, for mock clinical DBS samples, run on 1% agarose gel. The same samples were used as template DNA as in gel (B). All template DNA was diluted to 5-10ng/µl; 4µl was used as input for the multiplex drug resistance and *csp* PCR, 2µl was used as input for the *msp1* PCR, both to a final reaction volume of 50ul. 4µl of each PCR reaction was run on the gel. DBS = Dried Blood Spots. Neg = Negative control (Nuclease Free Water used as template).

The workflow was deployed on three sample sets: first, it was validated on laboratory parasite clones (3D7, Dd2, HB3, 7G8, GB4, KH1, and KH2) and mock clinical Dried Blood Spot (DBS) samples, referred to collectively as Validation samples (**Methods, Supplementary Figure 1**). Second, we performed prospective genomic surveillance on leucodepleted venous blood samples from clinical malaria samples collected at two sites in Ghana. Third, we retrospectively applied the workflow to a collection of dried blood spot samples collected in northern Ghana that had been used in another study. Nanopore sequencing was performed in multiplexed batches on an ONT MinION mk1b device using either kit 12 with R10.4 flow cells (Validation and leucodepleted venous blood samples) or kit 14 with R10.4.1 flow cells (Validation and clinical dried blood spot samples) (**Table 2**, **Methods**). For the Validation samples, no discrepancies were identified between the key antimalarial resistance markers genotyped in the assay and the expected genotypes for the laboratory clones tested. The lab isolate Dd2 was noted to contain both N86Y and N86F variants in *mdr1* due to having multiple copies of this gene, as previously observed (e.g. [75], discussed further in **Supplementary Notes**). For the two lab isolate mixtures, the consensus genotype assigned matched the expected majority clone – for example, C580Y in *kelch13*, which is associated with artemisinin partial resistance, was correctly genotyped in both the ‘pure’ KH2 isolate (known to possess that marker), and the mixture of KH2 : 3D7 at 80 : 20 ratios, respectively. However, *kelch13* was wild-type with consensus genotyping for the mixture with KH2 : 3D7 at 20 : 80 ratios, as expected. Sequence reads were also mapped to the full 3D7 reference genome and manually inspected using the Integrative Genomics Viewer (IGV) tool, confirming the mixed sample at expected positions. In the mock DBS samples, drug resistance calls were concordant with the expected genotypes for the parasite clone used (Dd2), even at the lowest parasitaemias tested (predicted 0.01% infected red blood cells (RBCs)), for which bands were no longer appreciated by gel electrophoresis (**Figure 1**).

**Table 2.** Nanopore data generation. All sequencing was performed using an ONT MinION mk1b device with real-time base calling using either High Accuracy (HAC) or Super Accurate (SUP) settings with *guppy*. Samples were multiplexed using ONT native barcoding and sequenced in batches. Batches A – F comprised clinical malaria samples collected in Ghana. Most batches had 22 clinical samples, except for C which had 15; all batches included 1 positive control (gDNA from one of the Dd2, HB3 or KH2 clones) and 1 negative control. Batch E included 15 samples that were repeats of samples included in batches A – D, to assess for assay consistency. In total, 109 unique clinical samples were sequenced (sample flow chart in **Figure 2**). All clinical samples were sequenced using ONT Q20 chemistry (kit 12, R10.4 flow cells). In addition, a ‘Validation’ (V) set of samples was sequenced (gel image shown in **Figure 1**), comprising laboratory isolates (3D7, Dd2, HB3, 7G8, GB4, KH1 and KH2 clones), 2 clone mixtures, 8 mock DBS samples in pairs at each parasitaemia of 10%, 1%, 0.1% and 0.01% infected RBCs, and a negative control. The Validation set was sequenced both using ONT Q20 (‘V12’ batch) and Q20+ chemistry (‘V14’ batch, with kit 14 and an R10.4.1 flow cell running at 400bps). The V12 and V14 Validation samples were identical. Total data generated from the clinical samples (batches A – F) and median data generated from Q20 chemistry (batches A – F and V12) are shown. Compared with Q20, we found that Q20+ chemistry produced more data with improved pore retention during sequencing (**Supplementary Figure 2**). DBS = Dried Blood Spots.

| Sample batch name | ONT chemistry used | Amplicon samples per run (of which clinical samples) | Run time | Data produced (GB) | Estimated bases (Gb) | Reads generated (M) | Bases called (real-time), pass (Gb) |
| --- | --- | --- | --- | --- | --- | --- | --- |
| A | Kit 12, R10.4 | 24 (22) | 6 hours | 33.04 | 1.66 | 1.26 | 1.25 |
| B | Kit 12, R10.4 | 24 (22) | 6 hours 40 mins | 33.52 | 1.55 | 1.71 | 1.26 |
| C | Kit 12, R10.4 | 15 (13) | 6 hours | 37.05 | 1.62 | 2.08 | 1.27 |
| D | Kit 12, R10.4 | 24 (22) | 8 hours | 36.47 | 1.71 | 1.77 | 1.29 |
| E | Kit 12, R10.4 | 24 (22, with 15 repeats) | 8 hours | 31.96 | 1.49 | 1.65 | 1.19 |
| F | Kit 12, R10.4 | 24 (22) | 8 hours | 43.31 | 1.76 | 2.68 | 1.34 |
| V12 | Kit 12, R10.4 | 17 (0) | 6 hours | 39.13 | 1.86 | 2 | 1.61 |
| V14 | Kit 14, R10.4.1 | 17 (0) | 6 hours | 52.11 | 2.76 | 3.47 | 2.57 |
| **Total clinical (A – F)** | **Kit 12, R10.4** | **109** | - | **215.35** | **9.79** | **11.15** | **7.6** |
| **Median Kit 12/ R10.4 (A – F, V12)** | **Kit 12, R10.4** | - | - | **36.47** | **1.66** | **1.77** | **1.27** |

The Validation samples were sequenced using both kit 12/ R10.4 flow cells and kit 14/ R10.4.1 flow cells. Relative to R10.4 flow cells, we observed improved flow cell performance over the course of sequencing using the R10.4.1 flow cells (expected Q20+ accuracy) (**Supplementary Figure 2**), with increased total data generated (52GB vs 36GB), estimated bases (2.76GB vs 1.66GB), reads generated (3.47M vs 1.77M) and base-called pass bases (real-time super accurate *guppy* base calling; 2.57M vs 1.27M) (**Table 2**). These trends were consistent for multiple R10.4.1 flow cells.

### Clinical sample collection and study population characteristics

Prospective clinical sample collection took place in two locations in Ghana, one urban (LEKMA Hospital, Accra, on the coast) with perennial malaria transmission, and one rural (three sites in and around Navrongo, in the Upper East Region), where malaria transmission is highly seasonal (**Supplementary Figure 3**). Samples were collected August – September 2022 during the rainy, high transmission season. 142 patients with a positive *P. falciparum* Rapid Diagnostic Test (RDT) were recruited into the study; 42 from LEKMA Hospital and 100 from Navrongo (**Figure 2**). Samples were typically 0.5-2ml venous blood that underwent leucodepletion by centrifugation and Buffy coat removal (**Methods**). Samples from 33 patients were excluded from Nanopore sequencing, due to low parasitaemia (<20 parasites per 200 White Blood Cells, WBC), poor DNA yield post-extraction (<1ng/µl) or time constraints. This yielded a final sample set of 109 samples, 70 from Navrongo and 39 from Accra, which were taken forward for Nanopore sequencing and analysis.

**Figure 2.**
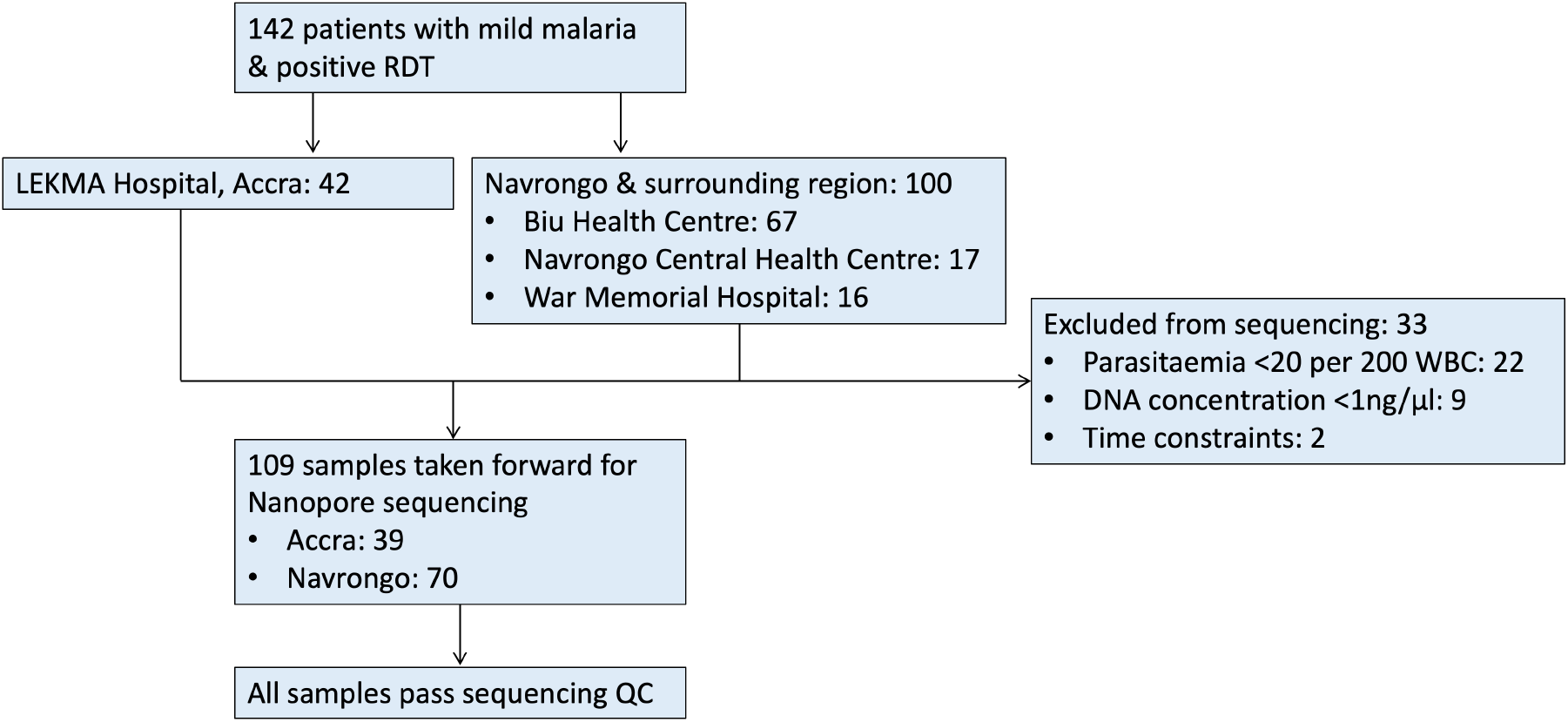
Study flow diagram. 142 patients with malaria symptoms and positive RDT were recruited into the study from LEKMA Hospital in Accra or three clinics in and around Navrongo in the Upper East Region. Samples were excluded if parasitaemia was <20 per 200 WBC, DNA concentration post-extraction <1ng/µl, or due to time constraints. (20 parasites/ 200 WBC corresponds approximately to 1,000 parasites per µl blood or 0.03% infected RBCs.) 109 samples were included in the study and taken forward for Nanopore sequencing. All samples, for all amplicons, produced >50x sequence coverage and were therefore included in downstream analyses. WBC = White Blood Cells.

For the 109 samples taken forward, median patient age was 12 years old (interquartile range (IQR) 5-22 years). There were 54 females and 51 males (4 unrecorded). Median parasite count was 864 (IQR: 243 – 1582) per 200 WBC; corresponding to approximately 43,000 parasites per µl blood (IQR: 12,000 – 79,000), or 1.4% infected red blood cells (RBCs) (IQR: 0.4% - 2.6%). The lowest parasitaemia included was 21 parasites/200 WBC, or approximately 1,000 parasites per µl blood (around 0.03% infected RBCs). For clinical samples collected in another study from 2015-2018 from mild malaria cases in Navrongo [22], a parasitaemia cut-off of 20 parasites/200 WBC would have included 72.3% of all samples (**Supplementary Figure 4**).

### Real-time multiplexed Nanopore sequencing on leucodepleted clinical samples

All of the 109 samples included were used for the multiplex drug resistance and *csp* PCR amplification assay, with encouraging gel electrophoresis results (**Supplementary Figure 5**). Using kit 12/ R10.4 flow cells, 6-8 hours of sequencing on the MinION mk1b in multiplexed batches of around 24 samples per flow cell produced a median of 34GB data, 1.62GB bases, 1.73M reads, and 1.26GB pass bases called per run (**Table 2**). Real-time base calling was performed using the Graphics Processing Unit (GPU) of a commercial gaming laptop and the resulting fastq files were used directly for downstream analysis.

The *nano-rave* pipeline can be run directly from the demultiplexed, base-called fastq files and folder organisation created in real-time during each ONT flow cell sequencing run, allowing rapid analysis. Median coverage across the amplicon targets was greater than 1000x per sample for all amplicons (range: 1552x median coverage for *csp* to 12141x for *dhfr*) (**Figure 3**), suggesting substantial scope for increased multiplexing to reduce costs. No amplicons from any sample in the 6-8 hour runs had <50x coverage, and therefore all samples were included in downstream genetic analyses; this suggested that lower parasitaemias and non-leucodepleted lower volume blood samples (such as DBS) could be used as sample input, which we subsequently confirmed (discussed below).

**Figure 3.**
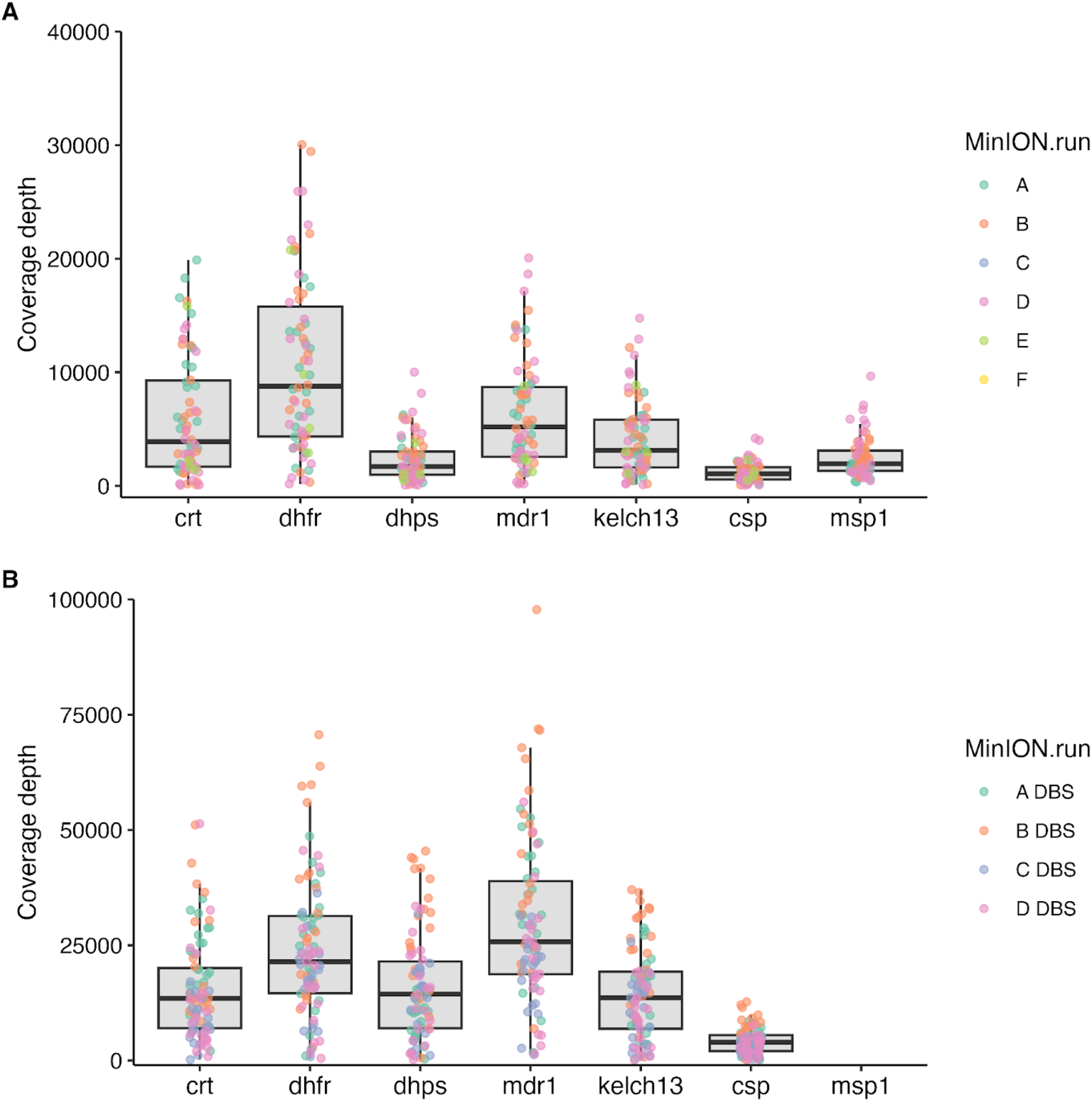
Coverage profile of amplicon targets. y-axis shows median number of reads covering each amplicon target per sample for each of the MinION runs for clinical samples. A) Leucodepleted venous blood samples, sequenced with kit 12 chemistry/ R10.4 flow cells (N=109 samples). Median coverage for the *crt*, *dhfr*, *dhps*, *mdr1*, *kelch13*, *csp* and *msp1* amplicons were, respectively: 8682, 12141, 2772, 8369, 5727, 1552, 1957. B) Dried Blood Spot (DBS) samples (N=87), sequenced with kit 14 chemistry/ R10.4.1 flow cells. In both figures, positive controls and sample duplicates were excluded. Coverage data were derived from BEDTools produced in the *nano-rave* pipeline. Note that the *msp1* PCR was only included in the leucodepleted venous blood samples from Navrongo (70/109), so coverage is only shown for these samples. Coverage was higher for the DBS samples however this may reflect using the newer ONT chemistry and flow cells.

To streamline the workflow and reduce informatic requirements, we aimed to genotype Single Nucleotide Polymorphisms (SNPs) using majority consensus calls, i.e. for genotypes from samples with mixed infections (more than one parasite clone present in the sample) to be based on the genotype of the most abundant clone. Three variant calling tools are currently available through the *nano-rave* pipeline: *medaka variant*, *medaka haploid* [76] and *freebayes* [77] (further information in **Supplementary Notes**). *Medaka haploid* was the fastest of these variant callers and felt to be the best suited for a haploid genome with Nanopore reads. For 14 samples, the workflow from PCR through to sequencing and variant calling was repeated to assess for assay consistency. No discrepancies were observed between the repeated samples using *medaka haploid* variant calling from *guppy* super accuracy or high accuracy base called reads, enabling these genotypes to be used for downstream analysis.

### Multiplexed Nanopore sequencing from clinical dried blood spot samples

Dried blood spots (DBS) are more readily collected than venous blood samples in resource-limited settings and are less invasive for most patient groups. The capacity to sequence directly from DBS samples therefore substantially extends the potential applicability of this workflow for malaria genomic surveillance in endemic countries. We tested the workflow using the multiplex drug resistance marker and *csp* PCR (with minor modifications to the PCR conditions, described in **Methods**) retrospectively on a set of 87 microscopy positive DBS samples collected from Navrongo in 2019. These samples were collected as part of ongoing surveillance work in the region, and already known to pass MalariaGEN quality control filters with Illumina whole genome sequencing (described in [78]). Median parasitaemia for the samples was 713 per 200 WBC (IQR 219 – 1882); the lowest parasitaemia sample had 2 parasites per 200 WBC (approximately 100 parasites per μl blood), i.e. close to the limit of microscopy positivity. As expected, lower parasitaemia samples had less *P. falciparum* and more human gDNA detected by qPCR (**Supplementary Figure 6**). Kit 14/ R10.4.1 flow cells were used in multiplexed batches of 24 samples per flow cell, run for 6-8 hours on a MinION mk1b device, with real-time super accurate base calling and genotyping using medaka-haploid in the *nano-rave* pipeline. Each flow cell run included a positive and negative control and a single sample was sequenced twice to compare between-run consistency.

Consistent with the mock DBS samples, bands were visible post-PCR by gel electrophoresis down to very low parasitaemias (**Supplementary Figure 7**). Amplicon coverage was high (**Figure 3B**); all amplicons had at least 50x coverage, including the samples with parasitaemias of <20 parasites per 200 WBC, and so were included in downstream analysis. Antimalarial resistance marker frequencies were consistent between the venous blood and DBS samples (**Table 3**, **Supplementary Figure 8).** The workflow was repeated twice for a single DBS sample, from PCR to sequencing, and again no discrepancies between repeats were identified. These data suggest that ONT can be used for amplicon sequencing of *P. falciparum* directly from DBS samples even at very low (but still microscopy positive) parasitaemias, without requiring a selective whole genome amplification step.

**Table 3.**
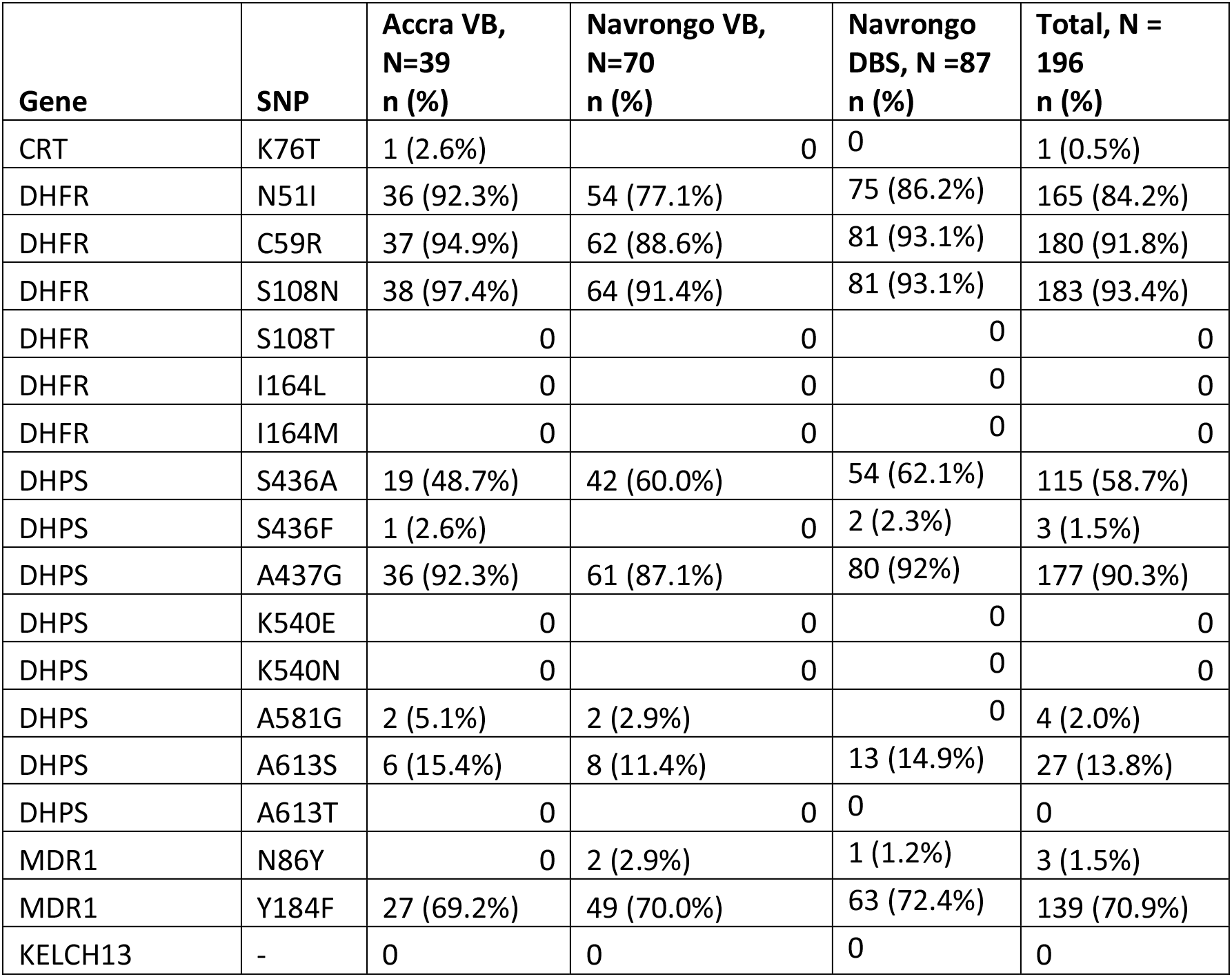
Antimalarial drug resistance genetic marker frequencies, for the combined 196 samples (109 leucodepleted venous blood and 87 dried blood spot samples). The table shows sample counts for the non-reference allele for each SNP, and non-reference allele frequency in brackets. Denominators are 39 venous blood samples from Accra, 70 venous blood samples from Navrongo, 87 dried blood spot samples from Navrongo, and 196 samples in total. For *kelch13*, all mutations were investigated and although five SNPs were identified, three were synonymous and the two non-synonymous changes were not known to be associated with artemisinin partial resistance – details in main text. VB = Venous Blood; DBS = Dried Blood Spot; SNP = Single Nucleotide Polymorphism; CRT = Chloroquine resistance transporter; DHFR = Dihydrofolate reductase; DHPS = Dihydropteroate synthase; MDR1 = Multidrug resistant protein 1.

| Gene | SNP | Accra VB,<br>N=39<br>n (%) | Navrongo VB,<br>N=70<br>n (%) | Navrongo<br>DBS, N =87<br>n (%) | Total, N =<br>196<br>n (%) |
| --- | --- | --- | --- | --- | --- |
| CRT | K76T | 1 (2.6%) | 0 | 0 | 1 (0.5%) |
| DHFR | N51I | 36 (92.3%) | 54 (77.1%) | 75 (86.2%) | 165 (84.2%) |
| DHFR | C59R | 37 (94.9%) | 62 (88.6%) | 81 (93.1%) | 180 (91.8%) |
| DHFR | S108N | 38 (97.4%) | 64 (91.4%) | 81 (93.1%) | 183 (93.4%) |
| DHFR | S108T | 0 | 0 | 0 | 0 |
| DHFR | I164L | 0 | 0 | 0 | 0 |
| DHFR | I164M | 0 | 0 | 0 | 0 |
| DHPS | S436A | 19 (48.7%) | 42 (60.0%) | 54 (62.1%) | 115 (58.7%) |
| DHPS | S436F | 1 (2.6%) | 0 | 2 (2.3%) | 3 (1.5%) |
| DHPS | A437G | 36 (92.3%) | 61 (87.1%) | 80 (92%) | 177 (90.3%) |
| DHPS | K540E | 0 | 0 | 0 | 0 |
| DHPS | K540N | 0 | 0 | 0 | 0 |
| DHPS | A581G | 2 (5.1%) | 2 (2.9%) | 0 | 4 (2.0%) |
| DHPS | A613S | 6 (15.4%) | 8 (11.4%) | 13 (14.9%) | 27 (13.8%) |
| DHPS | A613T | 0 | 0 | 0 | 0 |
| MDR1 | N86Y | 0 | 2 (2.9%) | 1 (1.2%) | 3 (1.5%) |
| MDR1 | Y184F | 27 (69.2%) | 49 (70.0%) | 63 (72.4%) | 139 (70.9%) |
| KELCH13 | - | 0 | 0 | 0 | 0 |

### Drug resistance marker frequencies

Antimalarial susceptibility was inferred from SNP genotypes using previously described inference rules [11] (**Table 3**, **Figure 4, Supplementary Notes**). Combining the prospective venous blood (n=109) and retrospective DBS samples (n=87), to yield a combined analysis set of 196 samples, we found that the vast majority (>99%) of samples were chloroquine susceptible, with only a single sample carrying the resistant *crt*-76T allele. There were high frequencies of resistant alleles to pyrimethamine (93.4% *dhfr*-108N) and sulfadoxine (90.3% *dhps*-437G). The majority genotype combination (for simplicity referred to as haplotype – caveats in Discussion) in *dhfr* was **<u>IRN</u>**I (83.2%) – the ‘triple mutant’, referring to amino acid positions 51, 59, 108 and 164 (wild-type = NCSI, mutant positions in bold and underlined). 8.7% of samples were N**<u>RN</u>**I – the ‘double mutant’ – or others (8.2%). No samples carried the highly resistant *dhfr*-164L allele. The two main *dhps* haplotypes identified were **<u>AG</u>**KAA (40.8%) and S**<u>G</u>**KAA (37.2%), i.e. (**<u>A</u>**/S)**<u>G</u>**KAA accounted for 78.1% of samples; this refers to *dhps* amino acid positions 436, 437, 540, 581 and 613 (fully susceptible = SAKAA; note that the 3D7 reference clone carries the resistant allele S**<u>G</u>**KAA). The most common *dhfr* and *dhps* haplotype combinations were *dhfr*-**<u>IRN</u>**I + *dhps*-**<u>AG</u>**KAA (35.7%) or *dhfr*-**<u>IRN</u>**I + *dhps*-S**<u>G</u>**KAA (30.6%). The haplotype combinations specifically associated with declining efficacy of SP-IPTp were not observed (defined as *dhfr*-51I, *dhfr*-59R, *dhfr*-108N + *dhps*-437G, *dhps*-540E + either *dhfr*-164L or *dhps*-581G or *dhps*-613S/T). However, all of these mutations except for *dhps*-540E were observed in this sample set. The prevalence of *dhps*-581G and *dhps*-613S mutations were 2.0% and 13.8%, respectively, and the *dhfr*-**<u>IRN</u>**I + *dhps*-**<u>AG</u>**KA**<u>S</u>** combination was present in 9.7% of samples. In *mdr1*, the frequency of the 86Y mutation was very low (3/196), while the 184F allele frequency was 70.9% (139/196).

**Figure 4.**
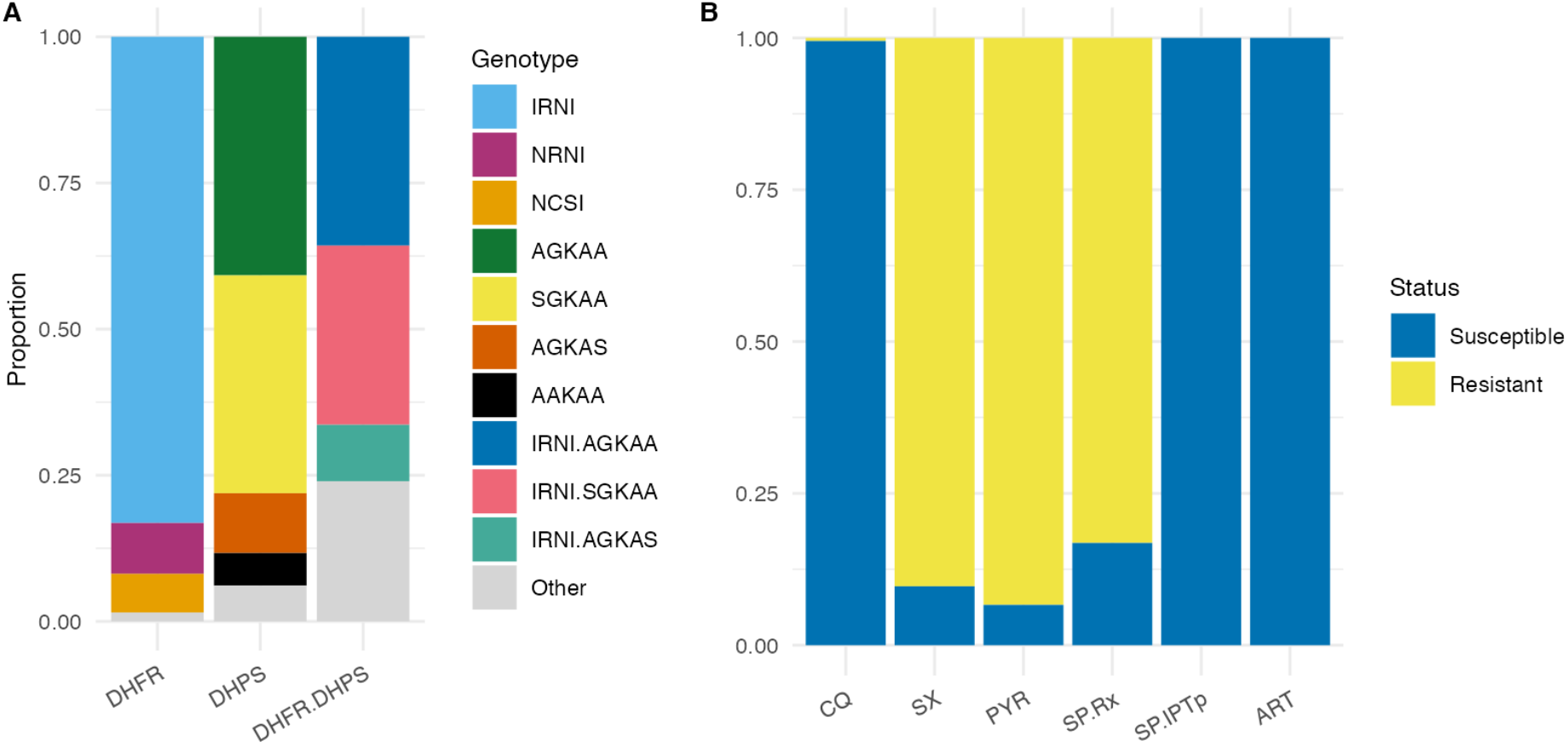
DHFR and DHPS haplotypes and inferred antimalarial resistance frequencies, for the combined 196 samples (109 leucodepleted venous blood and 87 dried blood spot samples). DHFR = Dihydrofolate reductase; DHPS = Dihydropteroate synthase; CQ = Chloroquine; SX = Sulfadoxine; PYR = Pyrimethamine; SP.Rx = Combination Sulfadoxine-Pyrimethamine (SP) as treatment for symptomatic malaria; SP.IPTp = Combination SP for intermittent preventive therapy in pregnancy; ART = Artemisinin. DHFR haplotypes refer to amino acid positions 51, 59, 108 and 164 (wild-type = NCSI). DHPS haplotypes refer to amino acid positions 436, 437, 540, 581 and 613 (fully susceptible = SAKAA). Inference rules for (**B**) are shown in **Supplementary Notes**. Note that for artemisinin, ‘resistance’ refers to artemisinin partial resistance (defined in main text). The leucodepleted venous blood and dried blood spot sample data are displayed separately in **Supplementary Figure 8**.

No mutations in *kelch13* were identified that have previously been associated with artemisinin resistance. Nine *kelch13* mutations were identified, five synonymous changes (two samples with C469C, and samples carrying T478T, A627A, and S649S), and four non-synonymous mutations: A578S, Q613H, N629Y, and V637I, all of which have previously been reported in Africa [79,80] and are not considered to be associated with artemisinin resistance (**Supplementary Notes**).

### Antigens and vaccine targets

We investigated SNP diversity in the C-Terminal Region (CTR) of *csp*, which is included in both the RTS,S/AS01 and R21-MM vaccines. Multiple SNP differences from the vaccine reference sequence were identified at high frequencies (>50% samples), resulting in amino acid changes such as S301N, K317E, E318(K/Q), N321K, and E357Q (**Figure 5, Supplementary Table 1**). The 301N mutation was present in 90% of samples. These SNP frequencies agreed very closely with whole genome sequence data using Illumina for *P. falciparum* in Ghana from the MalariaGEN Pf7 data resource [78] (**Supplementary Figures 9 – 10**). There was no evidence of population structure between the *csp*-CTR haplotypes present in Accra and Navrongo (**Figure 5B**). Overall, just 18 (9.2%) samples did not have any SNP mutations identified in the *csp*-CTR relative to the vaccine sequence. Parasites carrying an exact match to the RTS,S/AS01 or R21-MM *csp* haplotype were therefore a small minority of the parasite population in Ghana. However, our study did not assess whether the variants identified have any effect on vaccine efficacy.

**Figure 5.**
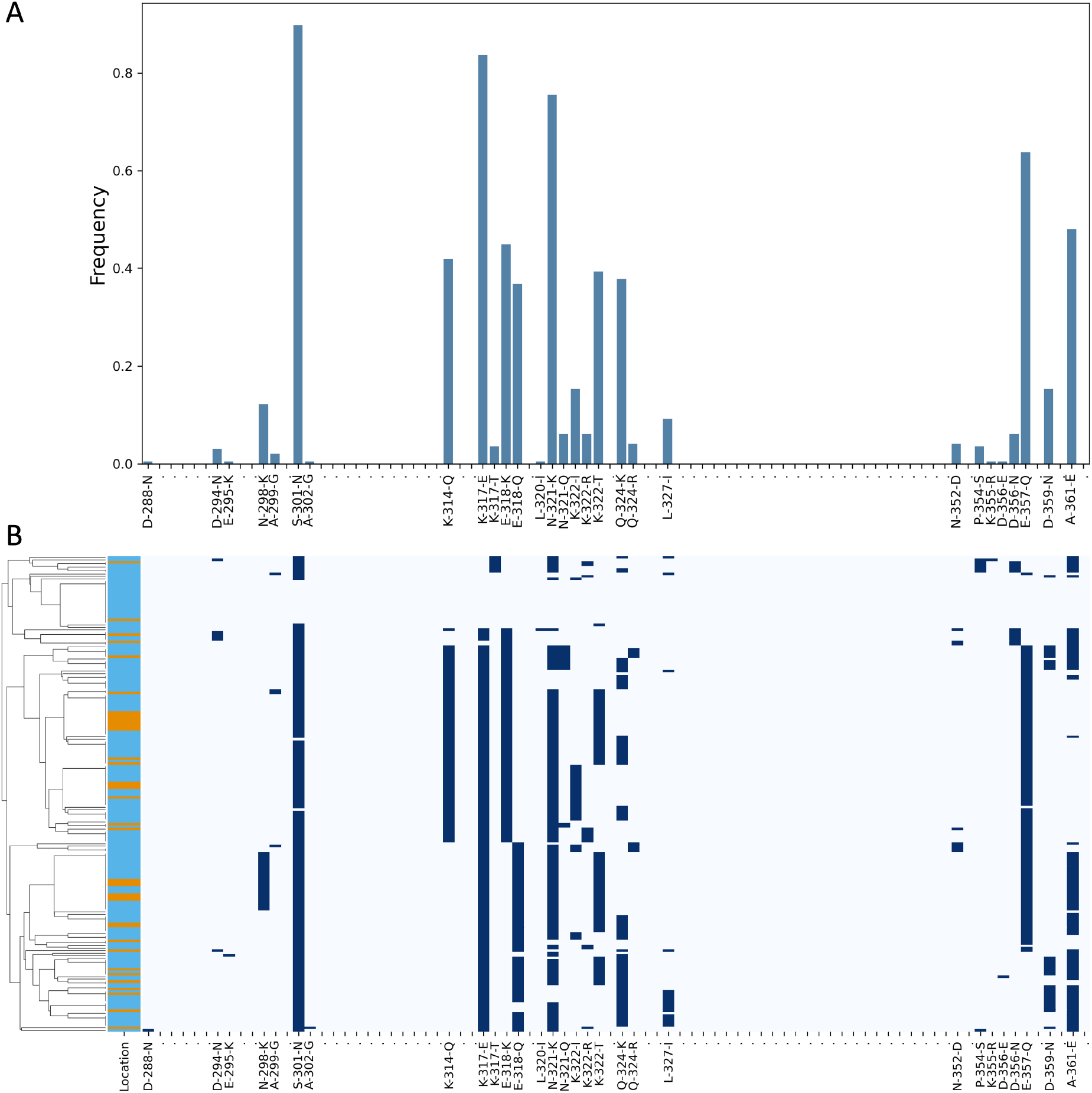
SNP frequencies in the *csp* C-Terminal Region. **(A)** Frequencies of SNPs along the C-Terminal Region (CTR) of *csp* identified from the Nanopore data (combining both the venous blood (n=109) and dried blood spot (n=87) samples, for total of 196 analysed samples), relative to the 3D7 reference sequence. Amino acid positions 288 to 362 (in the 3D7 reference) are displayed left to right ordered from N-terminal to C-terminal. Only variants with at least one sample carrying the non-reference allele are named in the x-axis. **(B)** Genotypes for the *csp* CTR for each sample in the study (N=196, rows), where dark blue = non-reference allele for that sample at that position and pale blue = the 3D7 reference sequence. Samples in **(B)** are grouped by haplotype similarity as represented by the dendrogram (left), with the colour bar indicating whether the sampling location was Accra (orange) or Navrongo (sky blue). Note that the *csp* CTR spans from amino acids 273 to 397 in the 3D7 reference, however no variants were identified in this cohort outside of the amino acids 288 to 362 displayed in the figure.

Lastly, we assessed Nanopore performance at producing accurate consensus full-length amplicon sequences, focusing on *csp* and *msp1* in the Validation samples. The *Amplicon_sorter* tool [81] was used to produce consensus sequences with a similarity threshold of 96% (details in **Methods**). Consensus sequences produced from PCR product in the expected ∼5Kb size range of the *msp1* amplicon (covering almost the entire *msp1* gene) had 100% base perfect mapping back to the reference sequences for all of the laboratory clones tested. For *csp*, base perfect consensus sequences were generated for the clones 3D7, Dd2, HB3, GB4, and KH2. Discrepancies were observed in the number of repeats in the Central Repeat region for two clones: in 7G8 there was a 12-bp deletion relative to the reference sequence (ATGCAAACCCAA). In KH1 there was a 24-bp insertion relative to the reference (GCAAACCCAAATGCAAACCCAAAT). It is possible that the reference sequences for these isolates are incorrect, or that the clones used for this experiment have altered during *in vitro* division relative to those used to produce the reference sequences.

## Discussion

We have performed multiplexed Nanopore sequencing and a rapid data analysis workflow from clinical *P. falciparum* malaria samples in two contrasting sites in Ghana. Genes of high clinical and public health importance were targeted for multiplexed PCR amplification, including drug resistance markers and the leading malaria vaccine and monoclonal antibody target, *csp*. The end-to-end workflow - including sample collection, DNA extraction, PCR, Nanopore sequencing and analysis of SNP markers - was performed at the field sites using standard molecular biology equipment and a handheld MinION. Real-time base calling and an informatics pipeline to call SNP variants were implemented on site using a commercially available gaming laptop. High bandwidth internet connectivity and high-performance computing clusters were not required. The key equipment could be transported between sites, located over 700km apart, including into a rural area around 200km from the nearest airport. The protocol was effective using both venous blood and dried blood spot samples, down to the lowest microscopy positive parasitaemias. A requirement for parasitaemia of >1 parasite per 200 WBC (approximately 100 parasites per μl blood) would be expected to capture the majority (>90%) of symptomatic malaria cases in Navrongo (**Supplementary Figure 4**). Though, altering the workflow conditions (e.g. increased sample multiplexing) would likely affect sensitivity. End-to-end time from sample collection to analysis output for a multiplexed batch of 24 samples was around 2 days, and this could be reduced further if needed. Given the high depth of sequence coverage achieved, increased sample multiplexing per MinION flow cell would very likely be successful. After relatively modest up-front hardware expenses (for example, a MinION mk1b is US$1,000 and a high-specification laptop may be around US$3,000), we estimate running costs of around US$35 per sample using this workflow with multiplexed batches of 96 samples. Higher levels of multiplexing and/or washing and re-using flow cells could reduce costs further.

Chloroquine resistance was highly prevalent (>80%) in Ghana in the early 2000s [82]. The data generated here indicate a trend towards increased chloroquine susceptibility, with nearly all samples genotyped as the wild-type *crt*-76K allele. This most likely reflects shifts in national treatment policy, as chloroquine was phased out due to resistance and replaced with alternative agents; ACT became the front-line antimalarial treatment in Ghana in 2005. Increasing chloroquine susceptibility has also been observed in Malawi [83]. In West Africa, the pattern is variable with very different chloroquine resistance rates observed even in nearby countries [78]. The high prevalence of *dhfr*-**<u>IRN</u>**I triple mutant (83%) parasites is broadly consistent with previous results from northern Ghana using short reads on a larger longitudinal sample set [22], in which the *dhfr*-**<u>IRN</u>**I triple mutant frequency was 67.9% in 2018. We also observed a high prevalence (78%) of *dhps*-(S/**<u>A</u>**)**<u>G</u>**KAA parasites. In [22], the frequency of *dhps*-(S/**<u>C</u>**/**<u>A</u>**)**<u>G</u>**KAA increased from 2.5% in 2009 to 78.2% in 2018; the ‘quintuple’ combination (*dhfr*-**<u>IRN</u>**I + *dhps*-**<u>AG</u>**KAA) was the dominant haplotype in 2018 at 76.6%. Although SP is no longer used as malaria treatment in Ghana, SP+AQ was introduced in 2016 as SMC targeting young children aged 3-59 months during the high transmission season in northern Ghana, and SP is also used as a prophylaxis in pregnancy (IPTp). Thus, there is continued parasite exposure to SP, which may be contributing to sustained and/or increasing resistant alleles in *dhfr* and *dhps*. While these mutations are associated with SP treatment failure for symptomatic malaria, there was no evidence of the high-level SP resistance marker *dhps*-540E that has been associated with reducing IPTp efficacy [25]. However, several other mutations such as *dhps*-581G and *dhps*-613S were identified. Given the use of SP for SMC and IPTp in northern Ghana, ongoing monitoring of *dhfr* and *dhps* will be an important component of malaria surveillance here. No *kelch13* mutations associated with artemisinin partial resistance were identified. Given the evolving situation with artemisinin partial resistance in East Africa, and the potentially devastating consequences of ACT treatment failure, molecular surveillance of markers for artemisinin and partner drug resistance remains critical in Africa. Thus, the successful end-to-end implementation of Nanopore sequencing in Ghana by a local team that works closely with the National Malaria Elimination Programme (NMEP) could compliment other platforms for malaria molecular surveillance, and enhance the rapid generation of data that is essential for monitoring antimalarial drug resistance to support NMEP decision-making.

The *csp* CTR harbours multiple SNPs relative to the reference sequence used in the RTS,S vaccine [84–90], and the more polymorphic regions correspond to T-cell epitopes [91]. The relationship between genetic diversity in *csp* and the efficacy of *csp*-based vaccines and monoclonal antibody therapies is incompletely understood, with conflicting findings for RTS,S (eg. [88] and [92,93]). While our study does not address this question, it demonstrates that Nanopore is an effective method for genotyping SNPs in the *csp* CTR as part of a multiplex surveillance panel, and identifies several high-frequency non-synonymous SNPs present in Ghana relative to the vaccine reference sequence. The SNP frequencies identified using ONT are very consistent with whole genome sequence data generated using Illumina in Ghana and West Africa more broadly [78]. Of note, Navrongo is one of the RTS,S/AS01 implementation sites and geographically close to the R21-MM vaccine trial site in Burkina Faso. Nanopore could therefore potentially be used for *csp* surveillance alongside monitoring drug resistance markers at little extra cost; future work could assess whether specific *csp* genetic variants – including any newly emerging variants as the RTS,S vaccine is rolled out – have an effect on vaccine efficacy.

Experience with SARS-CoV-2 has shown that prompt turnaround time is a key factor for genomic epidemiology to be useful in clinical and public health applications [48–50,94]. Results are available faster if sequencing can be performed in-country, without requiring the ethical, legal and logistical framework to transport samples outside of national borders. Sequencing workflows that can be implemented in endemic settings are essential to drive the decentralisation of genomics, to support its integration into clinical and public health applications, and to push for a more equitable distribution of global genomics capacity. Amplicon sequencing can be a pragmatic approach to malaria molecular surveillance and generate actionable data on parasite populations, including workflows based in endemic LMICs [11,12,103–107,95–102]. Advantages to amplicon sequencing include lower costs per sample, greater coverage and more reliable genotyping at loci of interest (such as drug resistance markers), less complex analysis pipelines, lower costs for storing and processing data due to lower data volumes, and faster turnaround times. However, whole genome sequencing (WGS) offers the potential to discover new variants without needing to design new amplicon panels, can generate deeper epidemiological insights such as through Identity By Descent (IBD) analyses, and can improve characterisation of certain complex forms of genetic variation such as structural rearrangements. Thus, a mixture of amplicon and WGS approaches can be beneficial, and ongoing technological developments and reduced sequencing costs may ultimately shift the balance in favour of WGS.

By developing and investing in sequencing capacity, the technical skills, experience and technology can potentially be applied to multiple high priority pathogens and emerging infection threats, maximising the impact of genomics in public health and strengthening global pathogen surveillance and health security [108]. The potential for an end-to-end turnaround time of 1-2 days also makes real-time *P. falciparum* sequencing a possible tool in clinical diagnostics, for example, to assess for *kelch13* mutations in patients with delayed parasite clearance or relapse following ACT. A key factor to ensure genomics can be deployed in endemic countries is to have stable supply chains for the procurement of laboratory consumables and sequencing hardware, and post-purchase technical support. Notwithstanding initiatives by the Africa CDC PGI to ease the supply chain barriers for sequencing reagents in general via centralized procurement and distribution to partner laboratories (sequencing hubs), improved delivery of reagents across Africa by Carramore, equitable access to reagents remains a challenge. It is unacceptable for sequencing equipment to be more expensive and/or more difficult to procure in endemic countries and this is an important barrier to be overcome. Moreover, although the workflow was relatively straightforward to implement, it nonetheless required a functioning molecular biology laboratory with access to a thermocycler for PCR, fridge and freezer, and gel electrophoresis. These are still significant barriers in many high endemicity regions. The workflow was also tested on symptomatic mild malaria cases; performance in asymptomatically infected people – an important reservoir – requires further study.

This study has several limitations. The *nano-rave* informatics workflow was designed to be streamlined and rapid, without requiring high performance computing clusters, based on majority SNP genotyping in amplicon targets. The workflow does not attempt to deconvolute mixed infections, making it unreliable to infer haplotypes (ie. genotypes shared within each clone). Future work could use full-length Nanopore reads to separate distinct haplotypes from mixed infections within samples. For calling SNP markers of drug resistance and SNP variation in *csp* for population-level surveillance, majority genotype calls will nonetheless provide useful information quickly. Copy number variation (CNV) was not assessed in this assay, such as amplifications in the drug resistance markers *mdr1* or *plasmepsin-2/3*. Calling CNV from PCR amplified products is inherently challenging with any sequencing platform, particularly for *mdr1*, which can have many break points causing multiple possible amplifications spanning large genomic regions [27]. Multiple extensions and/or modifications could be made to the PCR panel used in this study, depending on the specific use cases. For example, more of the *crt* gene could be included in the multiplex assay, given that variation along this gene has been associated with emerging partner drug resistance in Southeast Asia [10,109]; or adding *hrp2/3* targets to monitor for deletions. Future work can aim to increase the throughput of the assay, as our results demonstrate that higher levels of multiplexing are possible given the very high coverage achieved, and to determine whether the protocol can be used on ultra-low parasitaemia samples, such as microscopy negative samples and cases of asymptomatic infection. Further work can establish operational parasitaemia cut-offs for different sample types and sequencing approaches, using unselected clinical samples over a wide parasitaemia range. We also note that all genomic inferences of antimicrobial susceptibility carry a risk of failing to detect phenotypic resistance or predicting resistance that would not manifest *in vivo* in a given individual. This is true regardless of the sequencing platform used. Genomic predictions of antimalarial resistance in *P. falciparum* can be informative for monitoring markers of resistance at population-level, but linked phenotypic data remains essential to ensure that genetic markers are informative in specific populations.

Finally, while the decentralisation of genome sequencing offers many advantages, one potential downside could be a lack of standardisation, which may cause discrepancies between data from different studies and locations, making pooled analyses more challenging. One potential solution could be to establish a technical working group of scientists and public health experts active in malaria genomic surveillance using Nanopore, to agree on suggested best practices and processes for laboratory quality assurance. A commitment to open-access data sharing will be essential to ensure that locally produced data can be acted on quickly by the international community and integrated into larger analyses [110]. This would increase the breadth and depth of global malaria surveillance in the drive towards elimination.

## Methods

### Study setting

The study was based at two sites in Ghana with contrasting epidemiology: Ledzokuku Krowor Municipal Assembly (LEKMA) Hospital in Accra, and two satellite clinics in and around Navrongo and the War Memorial Hospital (WMH), in the Upper East Region near the northern border with Burkina Faso. LEKMA Hospital is in an urban setting near the coast where malaria is perennial, and represents a substantial burden of both inpatient and outpatient visits. Navrongo is a more rural setting, situated in a scrub-savannah ecological setting where malaria is strongly seasonal, with a high-transmission rainy season occurring around July – November. Sample collection for the venous blood samples took place August – September 2022 at both sites, so during the Navrongo high season. Sample collection for dried blood spots took place in Navrongo throughout 2019.

### Clinical sample collection and processing

This study incorporates samples collected under the governance of two separate studies. Samples from LEKMA Hospital were collected via the Emerging Genomic Selection and Antimalarial Drug Tolerance (EGSAT) study. Samples from Navrongo were collected via the Pan-African Malaria Genetic Epidemiology Network (PAMGEN) study. Both studies had approval from ethical review boards (ERB) for malaria parasite genomic sequencing research. In both sites, patients presenting with symptoms compatible with malaria were tested using Rapid Diagnostic Tests (RDTs) (OnSite Malaria Pf/Pan Ag Rapid Test by CTKBiotech, reference: R0113C). People positive for at least one of the Pf-specific antigen band (*hrp2/3*) and/or the pan-*Plasmodium* antigen band (LDH) were recruited with informed consent from the patient or their guardian. Around 2-5ml venous blood samples were collected, of which 0.5-4ml was typically available to use in this study.

Samples were transported daily, Monday – Friday, from LEKMA Hospital to the West African Centre for Cell Biology of Infectious Pathogens (WACCBIP), University of Ghana, and from the three Navrongo sites to the Navrongo Health Research Centre (NHRC) Research lab in Navrongo. Leucodepletion was performed by removing the Buffy coat layer following centrifugation, using the following steps: centrifuge blood sample in the EDTA tubes they were collected in at 500g for 5 minutes with no break, carefully remove the plasma and any visible Buffy layer, add approximately equal volume of PBS, repeat spin with same conditions, repeat PBS aspiration and any further visible Buffy coat plus the thinnest top layer of Red Blood Cells (RBCs, to maximise white blood cell removal). PBS was added to a final volume of 1-2ml, samples were transferred to 15ml falcon tubes and frozen in the −80 freezer until DNA extraction.

Samples from Navrongo included a prospective collection of leucodepleted venous blood (collected in 2022) and a retrospective selection of dried blood spot (DBS) samples, originally collected in 2019. The 87 DBS samples were randomly selected from a larger collection that had passed MalariaGEN quality control (QC) filtering for Illumina sequencing, with the added requirement for parasitaemia to be microscopy positive i.e. ≥1 parasite per 200 WBC by thick film microscopy.

### Mock clinical samples

Mock Dried Blood Spot (DBS) samples were produced by combining human whole blood ordered from Cambridge Bioscience Ltd with red blood cells (RBCs) infected with *P. falciparum* (Dd2 clone) cultured *in vitro*, and blotting 50µl onto Whatman 3M cards. *P. falciparum in vitro* culture was performed at the Wellcome Sanger Institute as described in [111]. Final haematocrit of the cultured parasite – whole blood mixtures was 35%. The volume of parasitised RBCs added to human whole blood was varied to produce an approximate final parasitaemia of 10%, 1%, 0.1% and 0.01% infected RBCs. The expected linear relationship between parasitaemia and the ratio of parasite to human DNA present in the mock DBS samples following DNA extraction was confirmed by quantitative PCR (qPCR) using probes targeting conserved regions of the *P. falciparum* and human genomes (**Supplementary Notes**).

### DNA extraction and quantification

Four methods for DNA extraction were used. For 87/109 of the prospectively collected venous blood samples, DNA extraction was performed using the New England Labs Monarch^®^ High Molecular Weight (HMW) DNA Extraction Kit for Cells & Blood (T3050) according to the manufacture’s protocol. 22/109 of the prospectively collected venous blood samples were extracted using the QIAmp® DNA Blood Mini Kit (51106) according to manufactures instructions with minor modifications detailed in Supplementary Notes. For the mock DBS samples, DNA extraction was performed using the QIAmp DNA Investigator Kit (56504), and the protocol was adapted from the ‘Isolation of Total DNA from FTA and Guthrie Cards’ with minor modifications detailed in Supplementary Notes. Finally, for the clinical DBS samples, DBS were transferred from Ghana to the Wellcome Sanger Institute and DNA was extracted using the QIAamp Investigator Biorobot kit on the Qiagen Biorobot Universal instrument using a custom protocol, described in Supplementary Notes.

DNA was quantified using ThermoFisher Scientific Qubit 2.0 Fluorometer with Qubit™ dsDNA high sensitivity (Q32854) and Qubit™ dsDNA broad range kits (Q32853), as per manufacturer’s instructions.

### Primer design and PCR amplification

Primers were designed using the *primer3* software [112–114]. Primer regions were selected based on sequence conservation after aligning target genes in *P. falciparum* from the reference genomes produced in [115]. Primer compatibility for multiplexing was assessed *in silico* using the Thermo Fisher Multiple Primer Analyzer (https://www.thermofisher.com/de/de/home/brands/thermo-scientific/molecular-biology/molecular-biology-learning-center/molecular-biology-resource-library/thermo-scientific-web-tools/multiple-primer-analyzer.html). Multiple iterations of primer combinations were tested and assessed by gel electrophoresis to identify the most robust combinations (producing the brightest bands down to the lowest parasitaemias with mock clinical DBS, and with minimal non-specific bands). Multiple iterations of PCR optimisation were undertaken to yield the final reaction conditions used.

All of the samples described in this study underwent multiplex drug resistance and *csp* amplification using the Thermo Fisher Platinum™ *Pfx* DNA Polymerase (11708039), with reaction conditions shown in **Supplementary Notes**. The Platinum™ *Pfx* DNA Polymerase enzyme has been discontinued by the manufacturer. We have found that the Kapa HiFi polymerase produces comparable results using the same primers. The *msp1* PCR was performed using Promega long-range GoTaq® Polymerase (M4021), reaction conditions in **Supplementary Notes**.

After PCR, a subset of samples from each 96-well plate, always including the positive and negative controls for that plate, were inspected by gel electrophoresis to ensure the PCRs had been successful (with blank negative controls) before proceeding to Nanopore sequencing. 2-4µl of the drug resistance and *csp* multiplex PCR was run for 45-90 minutes on a 2% agarose gel at 100V. 2-4µl *msp1* PCR was run for 45-90 minutes on a 1% agarose gel at 100V. PCRs were extracted and purified using the Qiagen MinElute PCR Purification kit (28004). The full volume of both the multiplex and *msp1* PCRs for each sample were combined at this stage, each being added to the same extraction column such that each sample yielded a single eluted solution including both PCR reactions. Samples were eluted in 100µl Elution Buffer. Post-extraction DNA quantification was performed using Qubit as described above.

### Nanopore library preparation and sequencing

For the prospective leucodepelted venous blood samples (n=109), library preparation was carried out using ONT kit SQK-NBD112.24 following the ‘ligation sequencing amplicons – native barcoding’ protocol. Manufacturer instructions were followed, except: Blunt/TA ligase Master Mix was substituted with NEB Quick T4 DNA Ligase and NEBNext Quick Ligation Reaction Buffer (5X) in the ‘native barcoding ligation’ (step 5) for three of the clinical sample libraries (‘D’, ‘E’ and ‘F’), due to running out of the Blunt/TA ligase Master Mix during field work without ready access to replacements. We did not observe any drop in yield for the libraries that used NEB Quick T4 DNA Ligase compared with Blunt/TA ligase Master Mix. Additional nuclease-free water was added to ensure a final volume of 20µl. For the negative controls, Nuclease Free Water was added to the same PCR reaction mixes and were taken through the full workflow including PCR, extraction and Nanopore library prep. Five batches of 24 and one of 15 samples were sequenced in six MinION runs, each with a fresh R10.4 flow cell; this included technical replicates for internal quality assessment. The MinION runs with clinical samples are referred to by the letters A to F in the main text. Every run included 1 positive and 1 negative control.

For the retrospectively sequenced batch of dried blood spot samples (n=87), sequencing was performed using ONT kit SQK-NBD114.24 following the ‘ligation sequencing amplicons – native barcoding’ protocol as per manufacturer’s instructions. Four batches of 24 samples were sequenced in four MinION runs, each with a fresh R10.4.1 flow cell; as with the venous blood samples, each run comprised 22 clinical samples, a negative control and a positive control. A single sample went through PCR and sequencing twice to assess for reproducibility of results.

The ‘Validation’ sample set of laboratory isolates and mock DBS samples was sequenced both with Q20 chemistry (same kit as above, SQK-NBD112.24) and with Q20+ chemistry (kit SQK-NBD114.24 on R10.4.1 flow cells) at 400bps.

### Hardware and workstation set-up

Sequencing, base calling and the real-time bioinformatic analysis were run from a commercial Dell gaming laptop with the following specifications: 11th Gen Intel Core Processor i7 (8 Core); 32GB (2x 16GB) DDR4, 3200MHz; GPU: NVIDIA GeForce RTX 3080 with 16GB GDDR6; 1TB M.2 Solid State Drive (SSD). During Nanopore sequencing, the laptop was connected to an Uninterruptible Power Supply (UPS) with surge protection. Additional fans were used to reduce laptop overheating.

### Bioinformatics

#### Real-time base calling and analysis using the *nano-rave* Nextflow pipeline

Base calling was done in real-time alongside sequencing using the MinKNOW software. We tested both High Accuracy (HAC) and Super Accurate (SUP) *guppy* base calling run via the laptop’s GPU. Analyses included in this study for the clinical samples were performed on HAC base-called reads. The resulting fastq files were processed through a custom Nextflow pipeline: *nano-rave* (Nanopore Rapid Analysis and Variant Explorer), run on the laptop using Debian as a Linux operating system for Windows. The *nano-rave* pipeline is available via GitHub at: https://github.com/sanger-pathogens/nano-rave. Briefly, following quality control (QC) metrics, sequence reads are mapped against 3D7 reference sequences for each of the amplicon target genes using *minimap2* [116]. Mapping to individual reference sequences for target genes, rather than to the whole genome, substantially reduces computational requirements for the workflow, allowing it to run at speed directly on a commercial laptop. .sam files are converted to .bam files using *samtools* [117]. Amplicon coverage data is generated using *BEDTools* [118]. There are three parametrised options available for variant calling: *medaka variant*, *medaka haploid* [76] and *freebayes* [77]. All three generate Variant Call Format (VCF) file outputs for each amplicon for each sample (ONT barcode). We tested *medaka variant* and *medaka haploid* on all clinical samples and used *medaka haploid* genotypes for downstream analyses described in the main text. VCF files were processed using custom R scripts to calculate SNP allele frequencies at key drug resistance loci. A cut-off of >50x coverage was applied for an amplicon to be included in the analysis; however, all amplicons for all samples in the study exceeded this cut-off. None of the negative controls included in this study generated directories that were >10MB in size, which was used as a parameterized cut-off in the *nano-rave* workflow; therefore, no negative controls were taken forward for real-time analysis.

#### Whole genome mapping and manual inspection

In addition to the real-time analysis performed on the laptop in Ghana outlined above, SUP base called reads were mapped genome-wide to the 3D7 reference genome using *minimap2* on the Wellcome Sanger Institute (WSI) High Performance Computing cluster (HPC). Read pile-ups for each amplicon locus were manually inspected using the Integrative Genomics Viewer (IGV) tool [119].

#### Consensus sequence generation for *csp* and *msp1*

Consensus sequences for *csp* and *msp1* were produced for the laboratory clones using *Amplicon_sorter* [81], a tool for building reference-free consensus sequences using ONT sequenced amplicons based on read similarity and length. Reads mapping to the 3D7 *csp* reference sequence were extracted and used as input for *Amplicon_sorter* using a similarity cut-off of 96% (the threshold for merging sequences to generate consensus). Because of high divergence from the 3D7 reference genome, the same approach could not be used for *msp1*; instead, reads in the expected size range (∼4700-5300bp) were pulled directly from the fastq files for consensus sequence building. For single laboratory clones, a threshold of 96% was used for consensus merging. For mixed isolates, this was increased to 98% to distinguish between the reference isolates used. The resulting consensus sequences were trimmed to include only the sequences within the primer sites and reverse complemented if needed. Consensus sequences were then mapped against the expected reference sequence using the Clustal Omega online tool.

### Ethics

The Navrongo samples were collected as part of the PAMGEN study, ethics approval ID: NHRC343, obtained from the Navrongo Health Research Centre (NHRC) Institutional Review Board. This includes both the prospectively sequenced leucodepleted venous blood samples (collected in 2022) and the dried blood spot samples (collected in 2019). The LEKMA Hospital samples (collected in 2022) were collected as part of the EGSAT study, ethics ID: ECBAS030/21-22, approved by the College of Basic and Applied Sciences Ethics Review Committee, University of Ghana. All study participants or their guardians gave informed consent.

## Author contributions

Conceptualization: WLH, LNA, DPK

Investigation: WLH, STG, EA, LNA, FNE, DKSJ, JMN, KB, AJRH, SN, SS, SA, EKA, CMM, VA, EAK, ED, MLP, SG

Formal analysis: WLH, LNA, STG, EA, KJ, RDP, JAG, AJRH

Software: OL, GvS, WB, WRS

Validation: WLH, STG, LNA, EA, KJ, RDP, JAG, AJRH

Data Curation: WLH, STG, LNA, EA, SN

Writing – original draft preparation: WLH, LNA

Writing – review and editing: WLH, LNA, STG, GAA, RDP, JAG, AJRH, CMM, SS, EA, EKA, ESA, DPK

Visualization: WLH, AJRH, JAG

Supervision: WLH, LNA

Project administration: WLH, LNA

Funding acquisition: DPK, LNA, GAA

## Acknowledgements

We thank Linzy Elton, Matt Dorman, Anna Kovalenko and Ian Goodfellow for their advice and suggestions on Nanopore sequencing. We thank Olivier Seret and Florent Lasalle for their contributions to creating the *nano-rave* analysis pipeline. We are grateful to staff at LEKMA Hospital, Accra, and the Navrongo Health Research Centre, Navrongo, who contributed to malaria sample collection. We are particularly grateful to Martin Awogbo, Kathrine Anuwe and Elizabeth Asobayire, Bismark Afari, Isaac Nyaaba and Charles Aforbiko for helping with the recruitment of patients. We thank the patients and their guardians who participated in the study.

This work is supported by Human Heredity and Health in Africa (H3Africa) grant H3A/18/002. H3Africa is managed by the Science for Africa Foundation (SFA Foundation) in partnership with Wellcome, NIH and AfSHG. The views expressed herein are those of the author(s) and not necessarily those of the SFA Foundation and her partners.

