## Supplementary materials for "Nanopore sequencing for real-time genomic surveillance of *Plasmodium falciparum*"

**Authors**

Sophia T. Girgis[1]*, Edem Adika[2]*, Felix E. Nenyewodey[3], Dodzi K. Senoo Jnr[3], Joyce M. Ngoi[2], Kukua Bandoh[2], Oliver Lorenz[1], Guus van de Steeg[1], Alexandria J. R. Harrott[1], Sebastian Nsoh[3], Kim Judge[1], Richard D. Pearson [1], Jacob Almagro-Garcia[1], Samirah Saiid[2], Solomon Atampah[2], Enock K. Amoako[2], Collins M. Morang'a[2], Victor Asoala[3], Elrmion S. Adjei[4], William Burden[1], William Roberts-Sengier[1], Eleanor Drury[1], Megan L. Pierce[1], Sónia Gonçalves[1], Gordon A. Awandare[2], Dominic P. Kwiatkowski[1], Lucas N. Amenga-Etego[2]**+, William L. Hamilton[1,5,6]**+

* These authors contributed equally

** These authors jointly supervised the work

**Affiliations**

1. Wellcome Sanger Institute, Wellcome Trust Genome Campus, Hinxton, CB10 1RQ, United Kingdom
2. West African Centre for Cell Biology of Infectious Pathogens (WACCBIP), College of Basic and Applied Sciences, University of Ghana, Legon, Ghana
3. Navrongo Health Research Centre (NHRC), Ghana Health Service, Navrongo, Upper East Region, Ghana
4. Ledzokuku Krowor Municipal Assembly (LEKMA) Hospital, Accra, Ghana
5. University of Cambridge, Department of Medicine, Cambridge Biomedical Campus, Hills Road, Cambridge CB2 0QQ, United Kingdom
6. Cambridge University Hospitals NHS Foundation Trust, Cambridge Biomedical Campus, Hills Road, Cambridge CB2 0QQ, United Kingdom

**+ Corresponding authors**

Dr. William L. Hamilton:

Dr. Lucas N. Amenga-Etego:

### Supplementary Figures

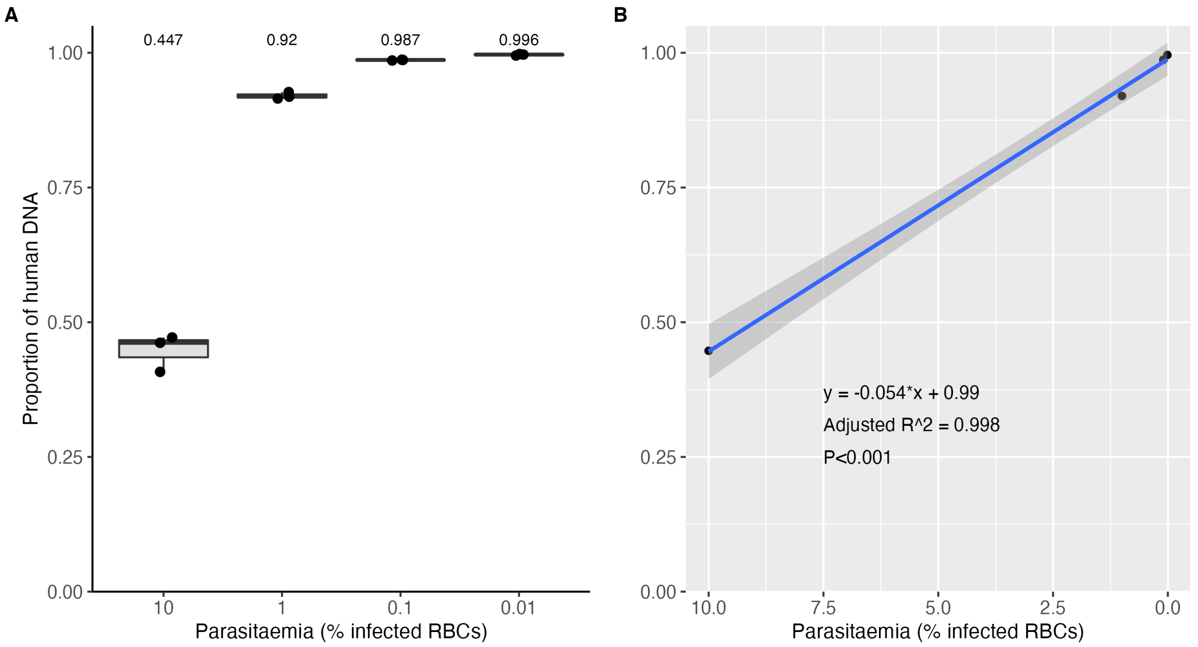


**Supplementary Figure 1.** Quantitative PCR (qPCR) results for mock clinical DBS samples. RBCs with *in vitro* cultured *P. falciparum* parasites were added to human whole blood in proportions estimated to yield parasitaemias of 10%, 1%, 0.1% and 0.01% infected RBCs. 50ul of these samples were blotted onto filter papers to produce mock DBS. Three DBS for each parasitaemia were DNA extracted (details in Methods). Extracted samples underwent qPCR for human and *P. falciparum* probes, with concentrations extrapolated from the standard curve. The y-axis for both plots shows the ratio of human to *P. falciparum* DNA concentrations, so values closer to 1 indicate more human DNA in proportion to parasite DNA and *vice versa* for values closer to zero. **A**) Boxplot of human DNA proportion for each parasitaemia (note non-linear x-axis). **B**) Linear regression model of median human DNA proportion vs parasitaemia. These results confirm a strong linear relationship between the relative abundance of *P. falciparum* DNA present in the mock DBS samples and the parasitaemia estimates, with high consistency between replicates. Further details on qPCR conditions are described in **Supplementary Notes**. DBS = Dried Blood Spot; RBC = Red Blood Cell.

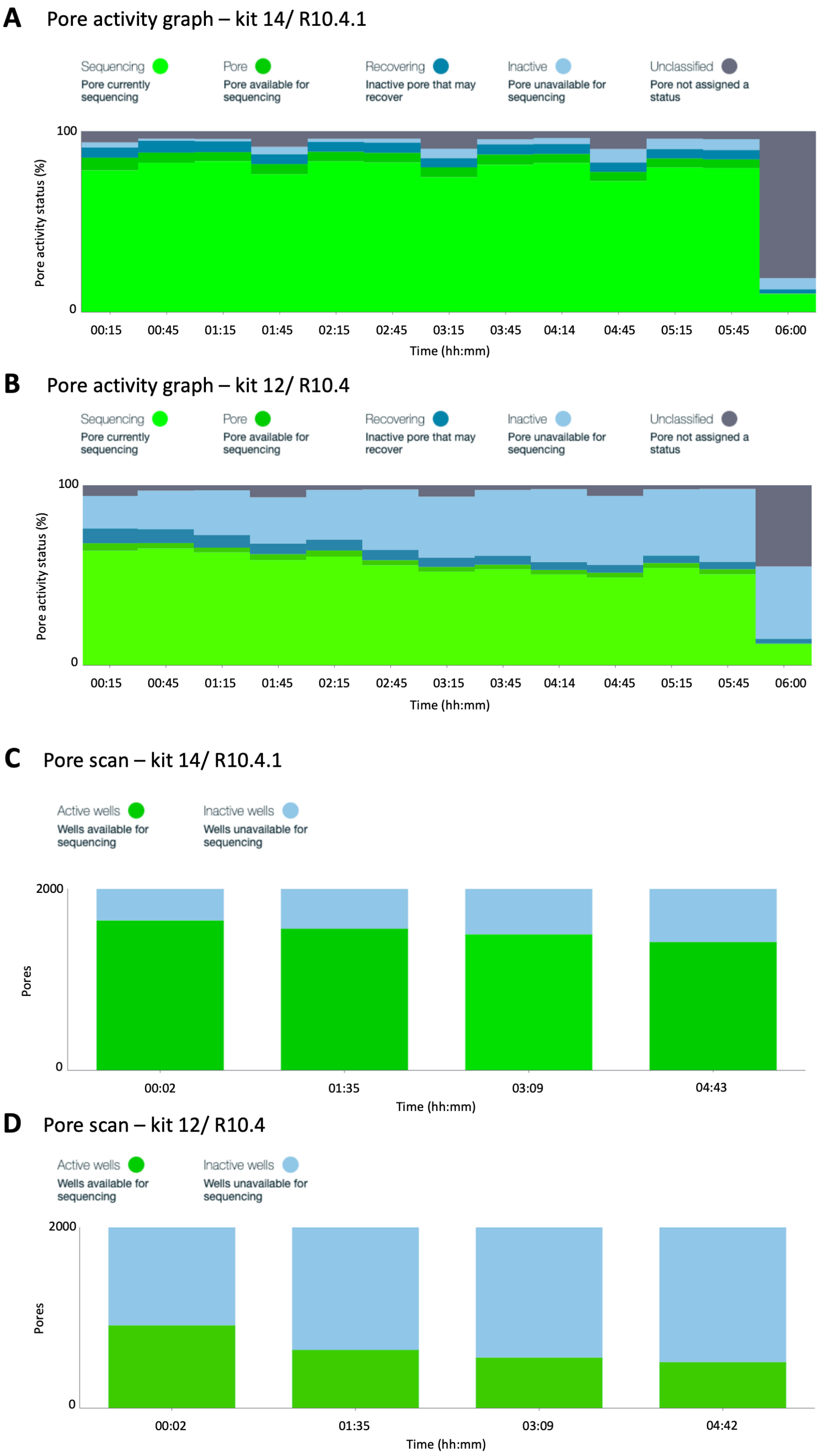
A
Pore activity graph – kit 14/ R10.4.1
Sequencing
Pore currently sequencing
Pore
Pore available for sequencing
Recovering
Inactive pore that may recover
Inactive
Pore unavailable for sequencing
Unclassified
Pore not assigned a status
100
0
Pore activity status (%)
00:15
00:45
01:15
01:45
02:15
02:45
03:15
03:45
04:14
04:45
05:15
05:45
06:00
Time (hh:mm)
B
Pore activity graph – kit 12/ R10.4
Sequencing
Pore currently sequencing
Pore
Pore available for sequencing
Recovering
Inactive pore that may recover
Inactive
Pore unavailable for sequencing
Unclassified
Pore not assigned a status
100
0
Pore activity status (%)
00:15
00:45
01:15
01:45
02:15
02:45
03:15
03:45
04:14
04:45
05:15
05:45
06:00
Time (hh:mm)
C
Pore scan – kit 14/ R10.4.1
Active wells
Wells available for sequencing
Inactive wells
Wells unavailable for sequencing
2000
0
Pores
00:02
01:35
03:09
04:43
Time (hh:mm)
D
Pore scan – kit 12/ R10.4
Active wells
Wells available for sequencing
Inactive wells
Wells unavailable for sequencing
2000
0
Pores
00:02
01:35
03:09
04:42
Time (hh:mm)

**Supplementary Figure 2 (previous page)**. Nanopore flow cell performance comparison: kit 14/ R10.4.1 flow cells (A, C) vs kit 12/ R10.4 flow cells (B, D). Showing screenshots from MinKNOW depicting pore activity (top; A, B) and pore scan results (bottom; C, D). Both sequencing runs used the multiplexed Validation sample set with native barcoding, and were run for 6 hours. Note the final data points for the pore activity plots is artefactual. Kit 14/ R10.4.1 is expected to produce Q20+ accuracy compared with Q20 for kit 12/ R10.4 flow cells. We have found that kit 14/ R10.4.1 flow cells produce higher and more sustained pore activity and almost no drop in pore activity after 6 hours of sequencing. This suggests that longer sequencing runs could be performed with ongoing high yield data generation, allowing for higher levels of multiplexing, and/or flow cell washes and re-use. Either of these approaches could increase throughput and reduce sequencing costs.

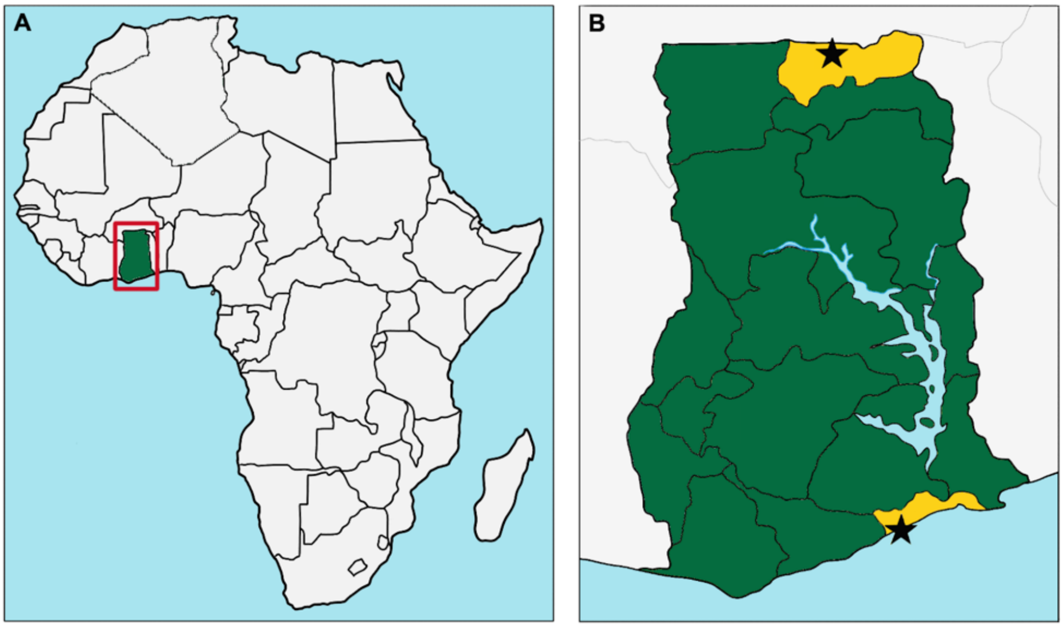


**Supplementary Figure 3**. Map showing location of Ghana in West Africa (A) and the two field sites within Ghana where the study was based (B), indicated by black stars: Lekma Hospital in Accra, near the coast, and Navrongo in the Upper East Region.

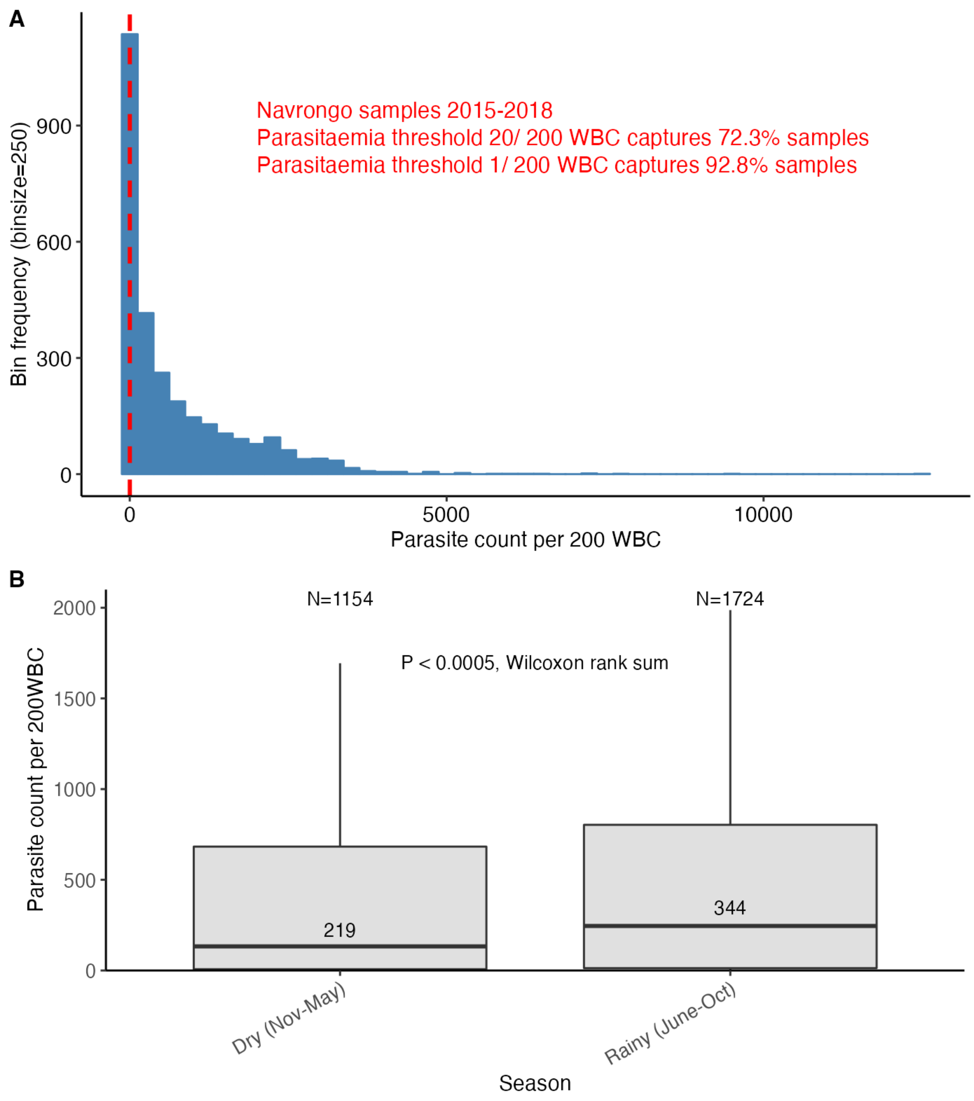


**Supplementary Figure 4.** Parasitaemia distribution for mild malaria cases sampled in the Upper East Region of northern Ghana, 2015-2018 (data gathered by Dr Lucas Amenga-Etego in previous studies). Cases were identified as having symptoms compatible with malaria and a positive Rapid Diagnostic Test (RDT). (**A**) Histogram of parasitaemias for all samples (n=2,878), indicating the minimum threshold used for the leucodepleted venous blood samples in this study of 20 parasites per 200 white blood cells (WBC) with dashed red line. Using this cut-off, 72.3% of all samples would have been included. For the Dried Blood Spot samples analysed in this study, we applied a cut-off of requiring microscopy positivity i.e. at least 1 parasite per 200 WBC. This would capture 92.8% of all samples from the Navrongo cohort shown in this figure, with similar values for both Dry and Rainy seasons. (**B**) Box plot of parasitaemias separated into samples collected during the Dry and Rainy seasons in Navrongo, defined roughly as November – May (n=1154) and June – October (n=1724), respectively. Transmission intensity is substantially higher during the Rainy season compared with the Dry season; consistent with this, a tendency for higher parasitaemias during the Rainy season is observed (median parasite count of 219 vs 344 in Dry vs Rainy seasons, respectively ($P$<0.0005, Wilcoxon rank sum test)). The cut-off of 20 parasites per 200 WBC would capture 69% and 74.6% of samples from the Dry and Rainy seasons, respectively. WBC = White Blood Cells.

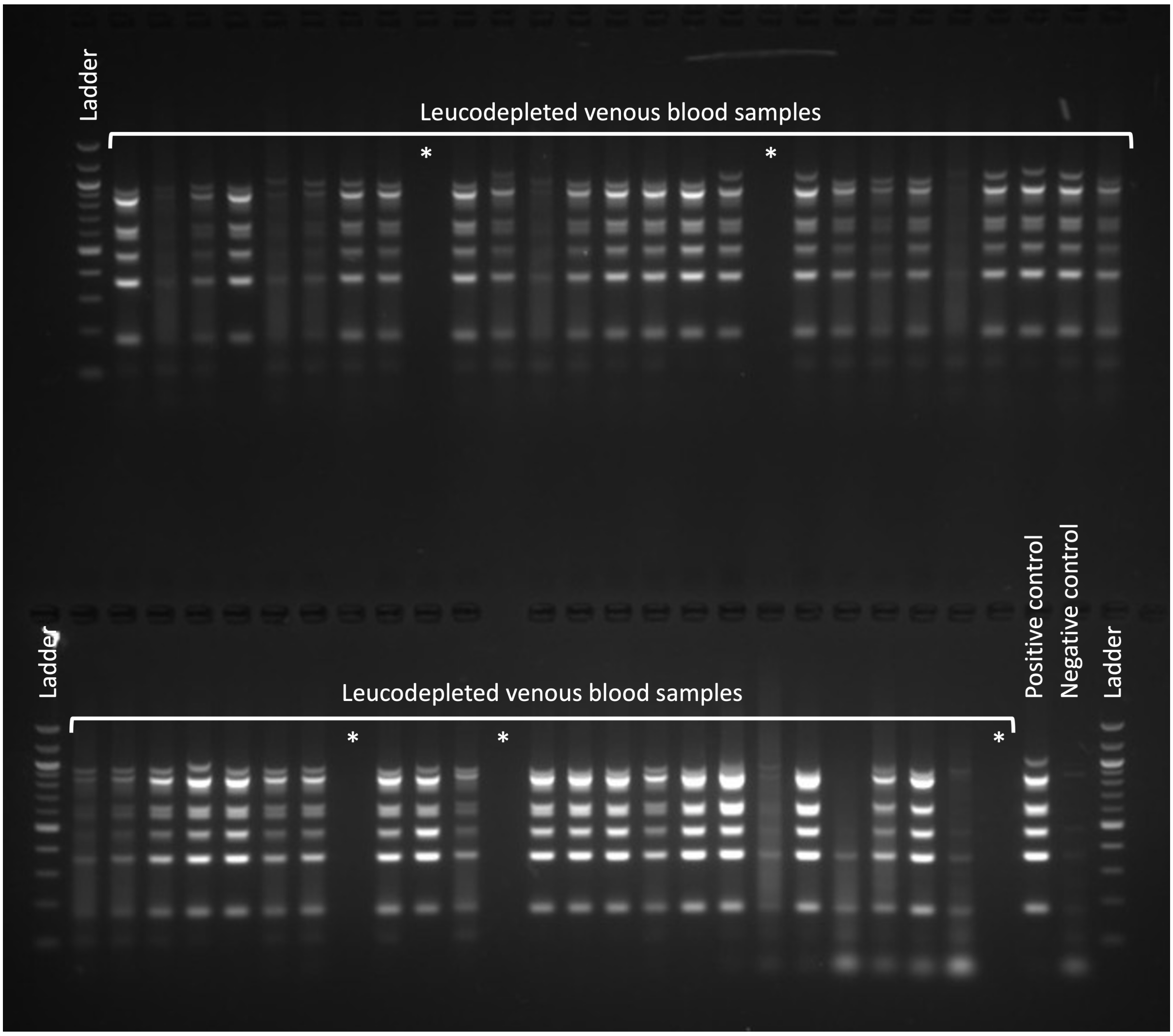


**Supplementary Figure 5.** Gel electrophoresis image of leucodepleted venous blood samples collected from patients with malaria at LEKMA Hospital, Accra, following multiplex drug resistance and *csp* PCR. Top: ladder in lane 1; samples in lanes 2 – 28 except for empty wells in lanes 10 and 19. Bottom: ladder in lanes 1 and 29; samples in lanes 2 – 25 except for empty wells in lanes 9, 13 and 26; positive control (pure *P. falciparum* gDNA) in lane 27; negative control (nuclease free water) in lane 28. Asterisks indicate empty lanes. Fragment size distribution follows the expected pattern, and is consistent with the laboratory and mock clinical isolates shown in **Figure 1**.

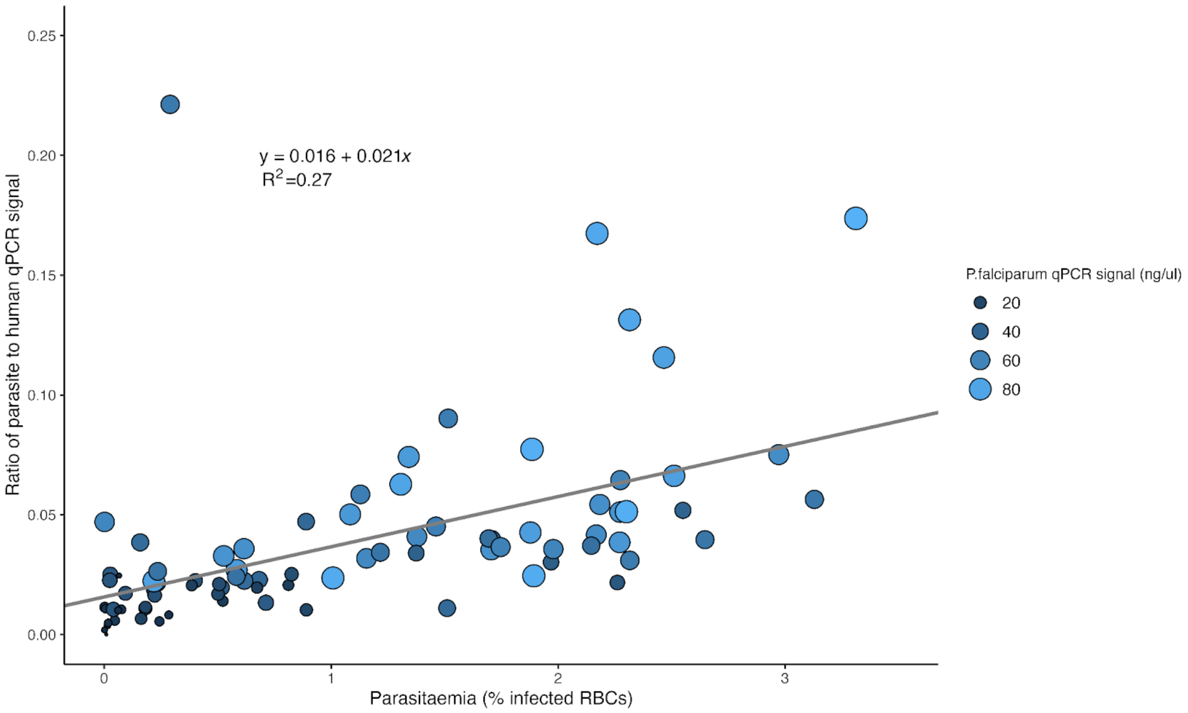


**Supplementary Figure 6.** The ratio of *P. falciparum* to human DNA measured by qPCR and the percentage parasitaemia (percentage of infected RBC) measured by microscopy, for the 87 dried blood spot samples collected from Navrongo and sequenced with ONT kit 14 chemistry/ R10.4.1 flow cells. Linear regression shows parasitaemia (%) is a significant predictor of the ratio of parasite to human DNA (*P*=2.13e-07), albeit with an $R^2$ of 0.27. qPCR methods are described in **Supplementary Notes**.

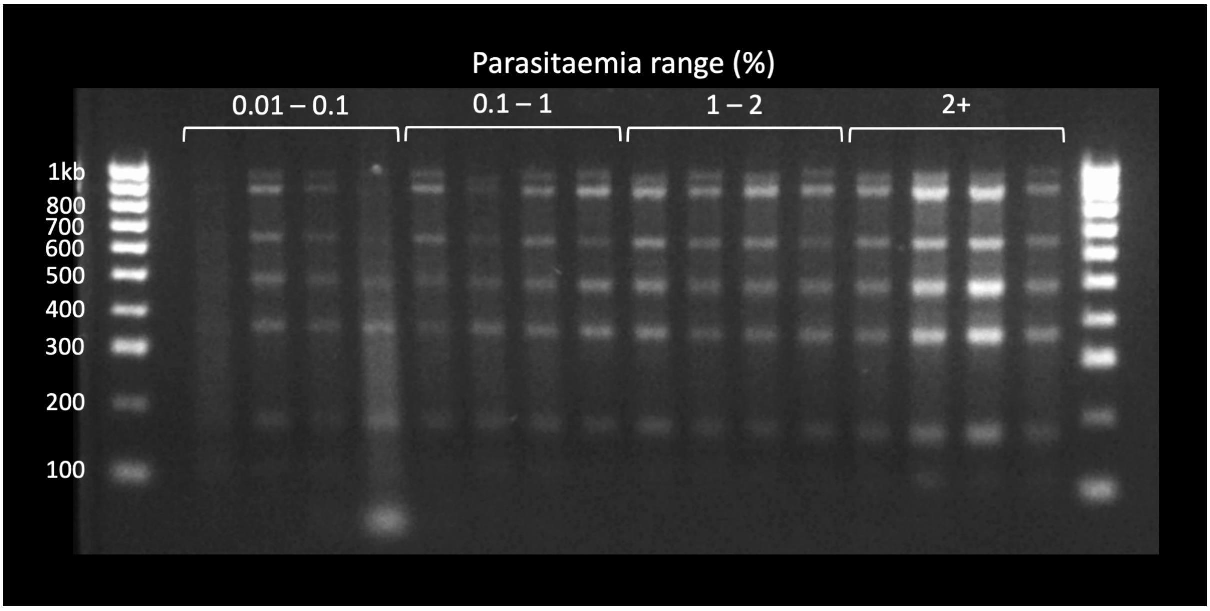


**Supplementary Figure 7.** Gel electrophoresis image of dried blood spot (DBS) samples collected from Navrongo, following multiplex drug resistance and *csp* PCR. Ladder in lane 1; samples organised left to right by increasing parasitaemias. Bands are visible at the expected sizes even for the lowest parasitaemias. Fragment size distribution follows the expected pattern, and is consistent with the laboratory and mock clinical isolates shown in **Figure 1** and the leucodepleted venous blood samples shown in **Supplementary Figure 5**.

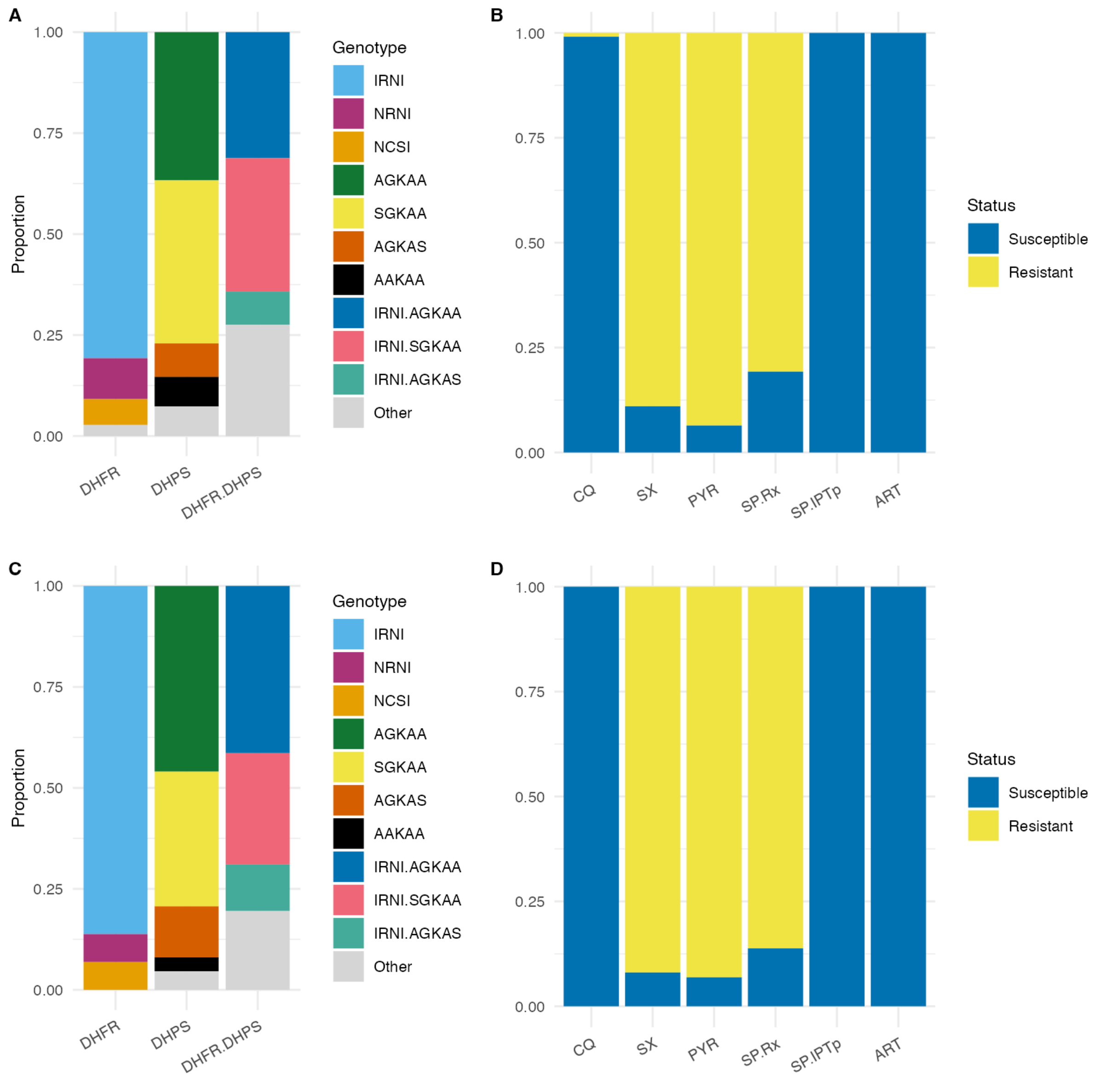


**Supplementary Figure 8.** DHFR and DHPS haplotypes (**A**) and inferred antimalarial susceptibility profile (**B**) for the 109 leucodepleted venous blood samples; and DHFR and DHPS haplotypes (**C**) and inferred antimalarial susceptibility profile (**D**) for the 87 dried blood spot samples. DHFR = Dihydrofolate reductase; DHPS = Dihydropteroate synthase; CQ = Chloroquine; SX = Sulfadoxine; PYR = Pyrimethamine; SP.Rx = Combination Sulfadoxine-Pyrimethamine (SP) as treatment for symptomatic malaria; SP.IPTp = Combination SP for intermittent preventive therapy in pregnancy; ART = Artemisinin. DHFR haplotypes refer to amino acid positions 51, 59, 108 and 164 (wild-type = NCSI). DHPS haplotypes refer to amino acid positions 436, 437, 540, 581 and 613 (fully susceptible = SAKAA). Inference rules for (**B**) and (**D**) are shown in **Supplementary Notes**. Note that for artemisinin, ‘resistance’ refers to artemisinin partial resistance (defined in main text). The leucodepleted venous blood and dried blood spot sample data are combined in **Figure 4** of main text.

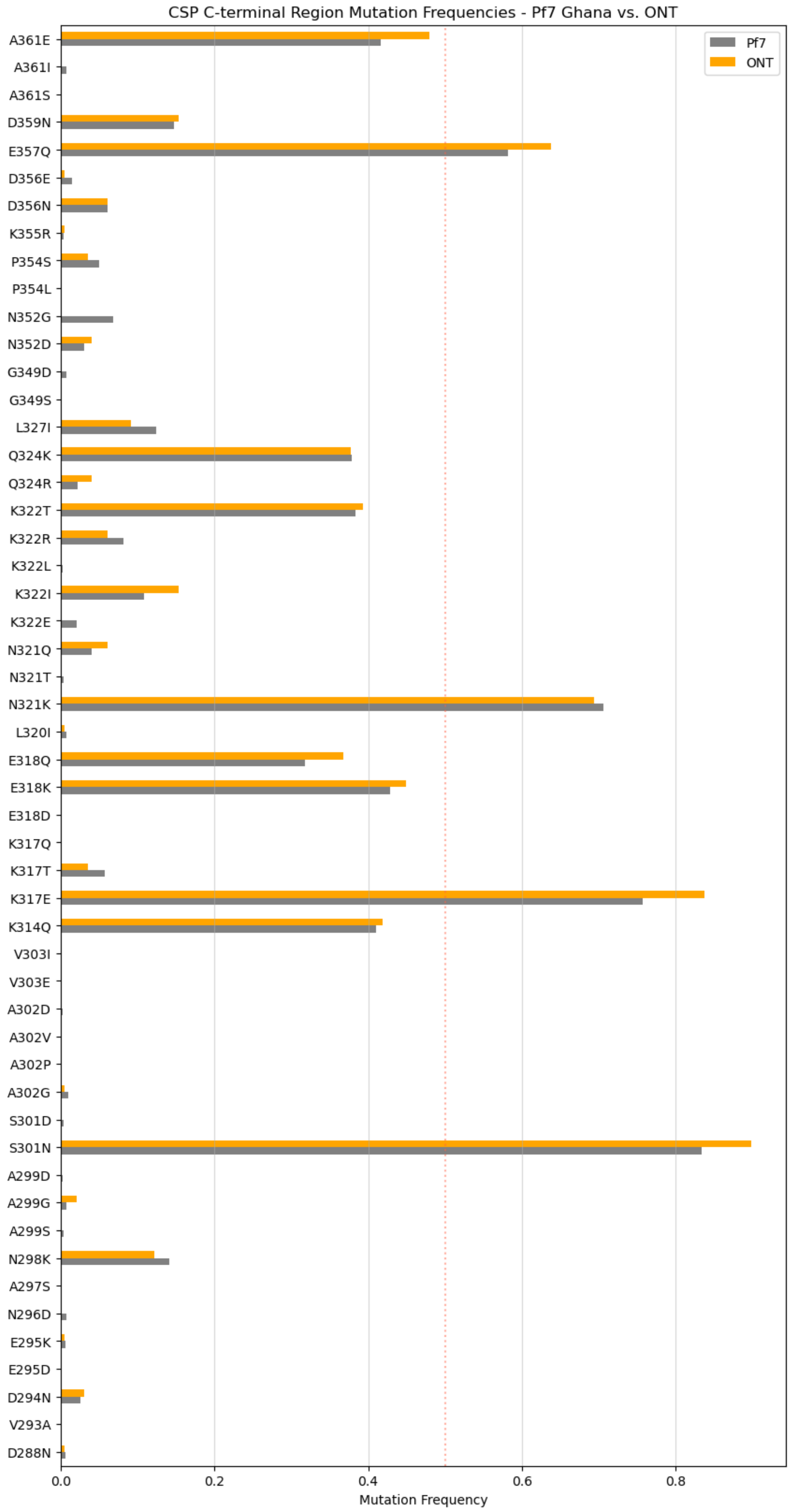
CSP C-terminal Region Mutation Frequencies - Pf7 Ghana vs. ONT
Pf7
ONT
A361E
A361I
A361S
D359N
E357Q
D356E
D356N
K355R
P354S
P354L
N352G
N352D
G349D
G349S
L327I
Q324K
Q324R
K322T
K322R
K322L
K322I
K322E
N321Q
N321T
N321K
L320I
E318Q
E318K
E318D
K317Q
K317T
K317E
K314Q
V303I
V303E
A302D
A302V
A302P
A302G
S301D
S301N
A299D
A299G
A299S
N298K
A297S
N296D
E295K
E295D
D294N
V293A
D288N
0.0
0.2
0.4
0.6
0.8
Mutation Frequency

**Supplementary Figure 9 (previous page).** Comparison of allele frequency estimates in Ghana for mutations in the C-terminal region of the csp gene. We show that frequency estimates for the ONT data gathered in this study (ONT, n=196) are very close to the estimates produced by the Ghanian samples of the MalariaGEN Pf7 dataset (WGS, n=1746). This analysis uses all C-terminal mutations observed in both datasets (selecting only samples from Ghana in Pf7) within clonal haplotypes (i.e., heterozygous mutation calls in Pf7 samples were discarded for frequency estimation). We also discarded Pf7 samples with missing data for the C-terminal haplotype. For a subset of mutations, ONT samples present a noticeable higher frequency estimate (e.g., S301N) but we confirmed with binomial tests (data not shown) that frequency differences could be explained by the variance introduced by the smaller ONT sample size (i.e., differences from ONT higher frequencies were not significant at p=0.05).

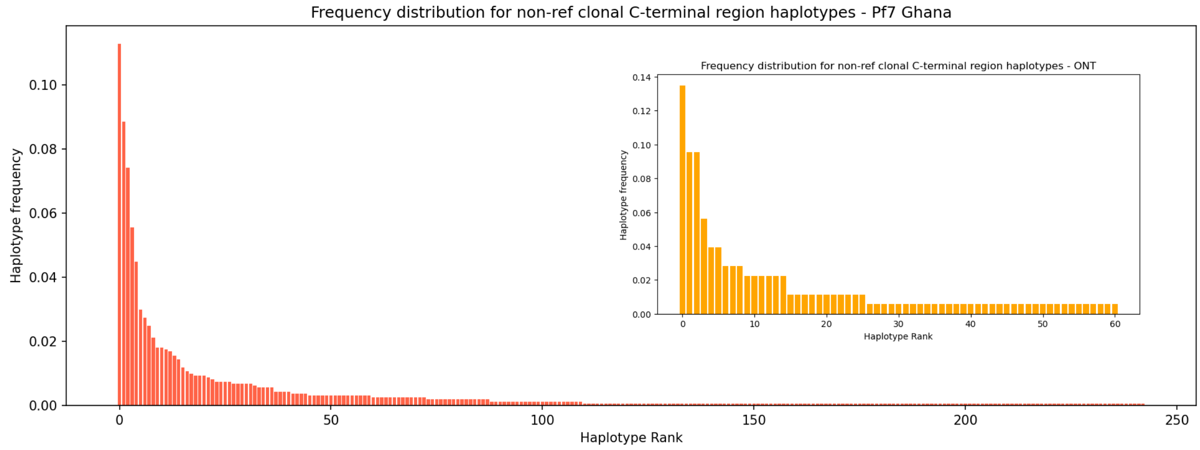


**Supplementary Figure 10.** Non-reference haplotype frequency distributions for the csp C-terminal region in samples from Ghana. We compare the ONT samples from this study (inset; ONT, n=178) with Ghanian samples in the Pf7 dataset (WGS, n=1604), after removing missing, heterozygous and reference haplotypes (i.e., haplotypes without any allele difference from the reference). Both distributions have a very similar shape, with a small set of high-frequency haplotypes that quickly decay into a long tail of minor ones. In addition, the first and third top-ranking haplotypes in both datasets are identical. This figure indicates that not only C-terminal mutations have very similar frequencies in both datasets (Supp. Figure 9) but that haplotype distribution and composition are also alike.

### Supplementary Tables

| DHFR haplotype | Haplotype count | Haplotype frequency | Haplotype % |
|---|---|---|---|
| IRNI | 163 | 0.83163265 | 83.2 |
| NRNI | 17 | 0.08673469 | 8.67 |
| NCSI | 13 | 0.06632653 | 6.63 |
| ICNI | 2 | 0.01020408 | 1.02 |
| NCNI | 1 | 0.00510204 | 0.51 |

**Supplementary Table 1.** DHFR haplotype frequencies. Showing haplotype frequencies (i.e. combinations of majority genotype calls) for the 196 samples analysed in the study (comprising 109 leucodepleted venous blood and 87 dried blood samples).

| DHPS haplotype | Haplotype count | Haplotype frequency | Haplotype % |
|---|---|---|---|
| AGKAA | 80 | 0.40816327 | 40.8 |
| SGKAA | 73 | 0.37244898 | 37.2 |
| AGKAS | 20 | 0.10204082 | 10.2 |
| AAKAA | 11 | 0.05612245 | 5.61 |
| SAKAA | 5 | 0.0255102 | 2.55 |
| AGKGS | 4 | 0.02040816 | 2.04 |
| SAKAS | 3 | 0.01530612 | 1.53 |

**Supplementary Table 2.** DHPS haplotype frequencies. Showing haplotype frequencies (i.e. combinations of majority genotype calls) for the 196 samples analysed in the study (comprising 109 leucodepleted venous blood and 87 dried blood samples).

| DHFR + DHPS haplotype | Haplotype count | Haplotype frequency | Haplotype % |
|---|---|---|---|
| dhfr-IRNI, dhps-AGKAA | 70 | 0.35714286 | 35.7 |
| dhfr-IRNI, dhps-SGKAA | 60 | 0.30612245 | 30.6 |
| dhfr-IRNI, dhps-AGKAS | 19 | 0.09693878 | 9.69 |
| dhfr-NRNI, dhps-AGKAA | 7 | 0.03571429 | 3.57 |
| dhfr-IRNI, dhps-AAKAA | 6 | 0.03061224 | 3.06 |
| dhfr-NRNI, dhps-SGKAA | 6 | 0.03061224 | 3.06 |
| dhfr-NCSI, dhps-SGKAA | 5 | 0.0255102 | 2.55 |
| dhfr-IRNI, dhps-SAKAA | 4 | 0.02040816 | 2.04 |
| dhfr-NRNI, dhps-AAKAA | 4 | 0.02040816 | 2.04 |
| dhfr-NCSI, dhps-AGKAA | 3 | 0.01530612 | 1.53 |
| dhfr-IRNI, dhps-AGKGS | 2 | 0.01020408 | 1.02 |
| dhfr-IRNI, dhps-SAKAS | 2 | 0.01020408 | 1.02 |
| dhfr-NCSI, dhps-AGKGS | 2 | 0.01020408 | 1.02 |
| dhfr-ICNI, dhps-AAKAA | 1 | 0.00510204 | 0.51 |

| | | | |
|---|---|---|---|
| dhfr-ICNI, dhps-SGKAA | 1 | 0.00510204 | 0.51 |
| dhfr-NCNI, dhps-SGKAA | 1 | 0.00510204 | 0.51 |
| dhfr-NCSI, dhps-AGKAS | 1 | 0.00510204 | 0.51 |
| dhfr-NCSI, dhps-SAKAA | 1 | 0.00510204 | 0.51 |
| dhfr-NCSI, dhps-SAKAS | 1 | 0.00510204 | 0.51 |

**Supplementary Table 3.** DHFR + DHPS combined haplotype frequencies. Showing haplotype frequencies (i.e. combinations of majority genotype calls) for the 196 samples analysed in the study (comprising 109 leucodepleted venous blood and 87 dried blood samples).

| SNP | Count | Frequency (%) |
|---|---|---|
| D288N | 1 | 0.51 |
| D294N | 6 | 3.06 |
| E295K | 1 | 0.51 |
| N298K | 24 | 12.2 |
| A299G | 4 | 2.04 |
| S301N | 176 | 89.8 |
| A302G | 1 | 0.51 |
| K314Q | 82 | 41.8 |
| K317E | 164 | 83.7 |
| K317T | 7 | 3.57 |
| E318K | 88 | 44.9 |
| E318Q | 72 | 36.7 |
| L320I | 1 | 0.51 |
| N321K | 148 | 75.5 |
| N321Q | 12 | 6.12 |
| K322I | 30 | 15.3 |
| K322R | 12 | 6.12 |
| K322T | 77 | 39.3 |
| Q324K | 74 | 37.8 |
| Q324R | 8 | 4.08 |
| L327I | 18 | 9.18 |
| N352D | 8 | 4.08 |
| P354S | 7 | 3.57 |
| K355R | 1 | 0.51 |
| D356E | 1 | 0.51 |
| D356N | 12 | 6.12 |
| E357Q | 125 | 63.8 |
| D359N | 30 | 15.3 |
| A361E | 94 | 48 |

**Supplementary Table 4.** List of Single Nucleotide Polymorphisms (SNPs) in the C-Terminal Region (CTR) of circumsporozoite protein (CSP). Ordered by amino acid position of discovered variants. Note the C-terminal domain in 3D7 can be considered as spanning amino acid positions 273 to 397. Nomenclature is based on differences from the amino acid position for the 3D7 reference sequence.

### Supplementary Notes

#### Study settings – further information

The study was based at two sites in Ghana with contrasting epidemiology: Ledzokuku Krowor Municipal Assembly (LEKMA) Hospital in Accra, and clinics based in and around Navrongo in the Upper East Region near the northern border with Burkina Faso. Sample collection in Navrongo took place at three sites: The Navrongo War Memorial Hospital (WMH) and Navrongo Central Clinic (NCC), within Navrongo town, and Biu Health Centre (BHC), around 30 km southwest of Navrongo. The NCC and BHC are community clinics, while WMH is the Municipal hospital with facilities for inpatient care. The Navrongo samples thus reflect community mild malaria cases that are RDT positive (baseline parasitaemia data for the region presented in Supplementary Figure 4). Samples from LEKMA hospital may represent a more selected patient group – both increased hospitalisation and prioritisation of which samples were collected from LEKMA may bias the cohort to higher parasitaemias, so direct comparison of parasitaemias between the Navrongo and LEKMA sites is confounded.

#### DNA extraction methods – further information

Four methods for DNA extraction were used. For 87/109 of the prospectively collected venous blood samples, DNA extraction was performed using the New England Labs Monarch® High Molecular Weight (HMW) DNA Extraction Kit for Cells & Blood (T3050) according to the manufacture's protocol. Part 1: erythrocyte lysis was conducted on frozen samples (-80°C) in 15 ml falcons with ~2 ml sample volume. After centrifugation, ~4-5 ml of supernatant was discarded. The pellet was dislodged, vortexed and transferred to a 2 ml Lo-Bind Eppendorf tube. A minimum of two 1xPBS washes were required and subsequent washes were carried out until the supernatant was clear. Part 2: leukocyte lysis: standard input volumes were required for all steps. Part 3: HMW gDNA binding and elution: isopropanol standard input volume; DNA was generally eluted in 110µl Elution Buffer (EB), though for a small number of samples where a large DNA pellet was visible, 210µl EB was used. Minor modifications were made to the protocol due to equipment availability; 1) samples were not kept on ice 2) all centrifuge steps were conducted at room temperature 3) samples underwent manual rotation as opposed to using a vertical rotating mixer 4) a heat-block replaced the incubator.

22/109 of the prospectively collected venous blood samples were extracted using the QIAmp® DNA Blood Mini Kit (51106) according to manufactures instructions with the following modifications: 1) 200 µl of frozen washed RBCs were thawed, and Protease was substituted with proteinase K. 2) Samples were incubated for 56°C for 30 minutes. 3) Samples were transferred into spin columns and centrifuged at 8000 rpm for 1 minute and 30 seconds. 4) Buffer AW1 was added and centrifuged at 8000 rpm for 1 minute and 30 seconds. 5) Buffer AW2 was added and centrifuged for at 13000 rpm for 3 minutes. 6)

An elution volume of 100µl was added to the spin columns and incubated at room temperature for 15 minutes. Extracted DNA was stored at -20°C.

For the mock DBS samples, DNA extraction was performed using the QIAmp DNA Investigator Kit (56504), and the protocol was adapted from the 'Isolation of Total DNA from FTA and Guthrie Cards'. Modifications to the protocol included adjusted quantities and an overnight incubation step: 1) 6 (1/8 inch) diameter punches placed into a 15 ml falcon tube, 2) 600 µl of Buffer ATL, 3) 60 µl of proteinase K, 4) Incubation at 56°C with shaking at ~600rpm for ~17 hours, 5) 600 µl of Buffer AL. Final steps were also modified to increase DNA concentration; 1) adjusted elution volume of 50 µl 2) spin column incubated for 5 minutes at room temperature 3) once centrifuged, eluate placed back into the spin column followed by a 5 minute room temperature incubation.

For the clinical DBS samples, DBS were transferred from Ghana to the Wellcome Sanger Institute. DNA was extracted using the QIAamp Investigator Biorobot kit on the Qiagen Biorobot Universal instrument using a custom protocol. For each DBS, 7 x 4mm discs were punched per well of a deepwell plate (puncher used: BSD600 Plus, BSD Robotics). A mastermix of buffer ATL and proteinase K was made and 660ul added per well (600ul ATL, 60ul proteinase K). The plates were incubated overnight at 56C for 17h, with shaking at 600rpm (Bioshake iQ, Q instruments). 600ul of lysate was transferred into a Qiagen S-block, which was loaded onto the Qiagen Biorobot Universal for extraction. On the Biorobot, a series of bind, wash and elution steps are performed. DNA is absorbed onto a silica membrane, and an integrated vacuum system draws lysates and buffers through, prior to eluting in 100ul nuclease free water. Key process modifications involved scaling up reaction volumes and increasing the elution incubation time to maximise DNA yield. Wash buffer volumes were increased accordingly, to minimise carryover of blood into the final eluate.

#### *nano-rave* pipeline and variant calling

An outline of the *nano-rave* pipeline is shown below:

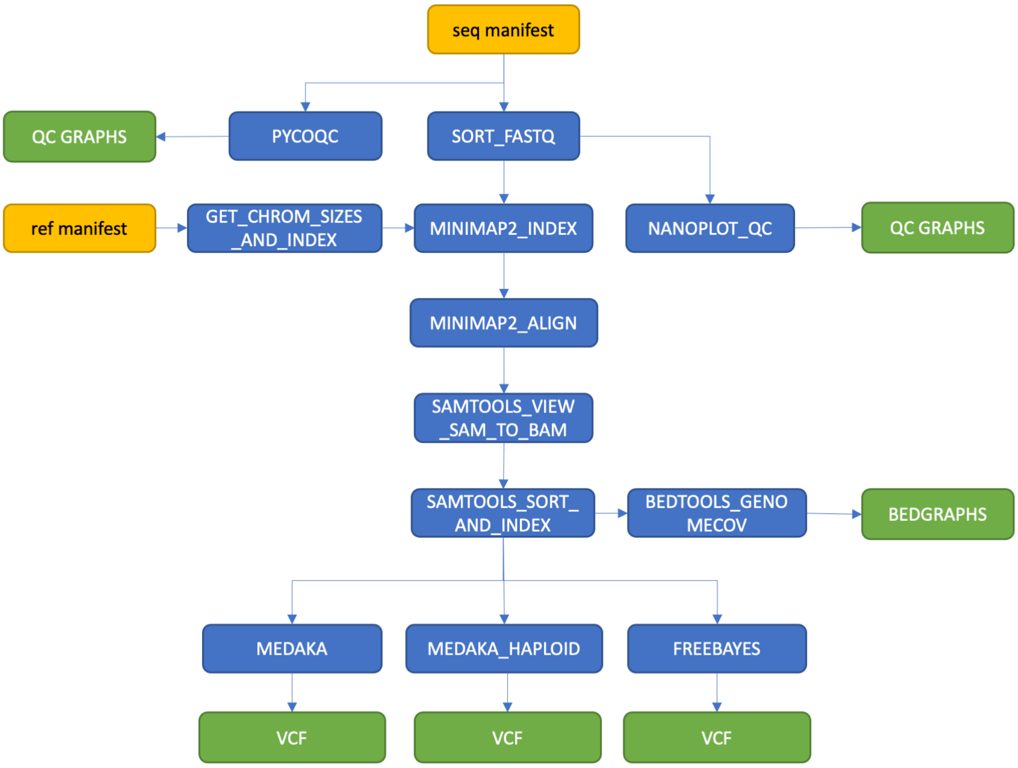


Full details of the pipeline including installation and user guides can be found at: https://github.com/sanger-pathogens/nano-rave.

We used *nano-rave* to align sequence reads against each amplicon reference (3D7) sequence, rather than against the full *P. falciparum* genome, to reduce computation requirements – although, this is not inherent to the pipeline and whole genome alignments could be performed. Three variant callers are currently available in the pipeline to produce VCF files. *Freebayes* is a Bayesian genetic variant detector designed for short read sequence data. *Medaka* uses neural networks to create consensus sequences and variant calls from Nanopore data. *Medaka_variant* was designed for diploid genomes and relatively lower coverage whole genome sequence data, and is being replaced by newer algorithms such as *Clair3*. *medaka_haploid_variant* was designed for calling variants in monoploid organisms, based on comparing the consensus sequence built in *medaka* against a reference sequence. We felt that *medaka_haploid_variant* (referred to as *medaka haploid*) was well suited to our use case, to generate majority genotype calls from the consensus sequence for each amplicon; 'heterozygous' positions (reflecting mixed infections) would therefore be genotyped by the majority allele present in that infection at that position.

The drug resistance SNPs genotyped using the *nano-rave* workflow with *medaka haploid* matched the expected positions for the reference isolates tested (3D7, Dd2, HB3, 7G8, GB4, KH1 and KH2). Aligned reads were also manually inspected using IGV. The clone mixtures tested in the Validation set (3D7 and KH2 clones in 80:20 or 20:80 ratios) confirmed this approach yielded the expected results, ie. genotypes were based on the majority clone in the mixture (see main text). We note that using this method, each locus strictly must be treated in isolation, making haplotype inference unreliable. This is because, depending on the proportion of clones present and their individual genotypes, the majority at one locus does not necessarily belong to the same haplotype as the majority at a different locus. Haplotype inference from deconvoluted mixed infections using Nanopore reads is an important area for future research. However, the majority SNP genotyping method applied here can nonetheless produce information useful for malaria surveillance rapidly, such as the prevalence of specific markers of antimalarial drug resistance.

On our laptop, the workflow for a multiplexed batch of 15-24 samples using *medaka variant* and *medaka haploid* variant calling took around 16-23 minutes, depending on the number of samples, amplicons and depth of coverage; *medaka haploid* tended to be slightly faster than *medaka variant*. *Freebayes* took greater than 45 minutes per multiplexed batch and so was not used further for real-time analysis of clinical samples. For 14 samples, the workflow from PCR to sequencing and variant calling was repeated to assess for assay consistency, using both *medaka variant* and *medaka haploid*. Using *medaka variant*, two discrepancies in drug resistance marker SNPs were identified. Both were samples with mixed infections in the discordant SNP as indicated by manual read inspection in IGV, with each sample in the discordant pair being assigned a different allele from the mixture in the two repeats. No discrepancies were identified in this sample set using *medaka haploid* genotype calls, and this variant caller was used for the downstream analyses described in main text.

We observed some inconsistency between sequencing repeats for the Dd2 clone in how position 86 of *mdr1* was genotyped using both *medaka variant* and *medaka haploid*. The wild-type (3D7 sequence) is 86-N, encoded by AAT; Dd2 contains both 86-Y and 86-F, encoded by TAT and TTT, respectively, in different copies of this gene within its genome. One of the mock DBS samples genotyped 86-F here, while the 7 mock DBS samples were 86-Y. Inspection of the read pile-ups in IGV suggested an approximately 50:50 ratio of these two SNPs, with 7 clones having slightly higher frequencies for 86-Y and the one 86-F clone having a slightly higher proportion of that SNP. This likely reflects chance variation in the workflow from PCR amplification to sequencing. Thus, there is potential for mixed infections (or heterozygosity in multi-copy genes, as in *mdr1*) to produce discrepancies in majority genotype calls, particularly if the proportion of each clone is similar and/or if the starting amount of parasite DNA is very low, as the effect of chance differences in PCR amplification during early cycles may be exaggerated. We note that this problem would also apply to Illumina-based amplicon sequencing methods.

#### PCR reaction conditions

Primer sequences for the combined drug resistance and *csp* PCR are shown below:

| **Primer name** | **Primer sequence** |
|---|---|
| crt F | TGTCTTGGTAAATGTGCTCA |
| crt R | AGTTGTGAGTTTCGGATGTT |
| dhfr F | GTTTTCGATATTTATGCCATATGTG |
| dhfr R | TGATAAACAACGGAACCTCC |
| dhps F | TTTGTTGAACCTAAACGTGC |
| dhps R | AACATTTTGATCATTCATGCAAT |
| mdr1 F | TGTGTTTGGTGTAATATTAAAGAACA |
| mdr1 R | ACATAAAGTCAAACGTGCATTT |

| | |
|---|---|
| kelch13 F | AAGCCTTGTTGAAAGAAGCA |
| kelch13 R | GGGAACTAATAAAGATGGGCC |
| csp F | TGGGAAACAGGAAAATTGGTAT |
| csp R | TACGACATTAAACACACTGGAA |

Primer sequences for the separate *msp1* PCR are shown below:

| **Primer name** | **Primer sequence** |
|---|---|
| msp1 F | AGAAGATGCAGTATTGACAGGT |
| msp1 R | GAACTGCAGAAAATACCATCGA |

The position of each amplicon within the respective target gene sequence, relative to 3D7 reference, is shown below:

| **gene** | **Amplicon start** | **Amplicon end** | **Amplicon size** |
|---|---|---|---|
| crt | 118 | 295 | 177 |
| dhfr | 22 | 512 | 490 |
| dhps | 1222 | 1863 | 641 |
| mdr1 | 228 | 591 | 363 |
| kelch13 | 1277 | 2145 | 868 |
| csp | 168 | 1143 | 975 |
| msp1 | 105 | 5099 | 4994 |

The reaction mixture for the combined drug resistance and *csp* PCR used for leucodepeleted venous blood samples is shown below for a single sample (these values would be multiplied accordingly to prepare master mixes). The concentration of gDNA used was typically 5-10ng/ul.

| **Reagent** | **1x for 50ul** |
|---|---|
| 10x Buffer | 6 |
| 10mM dNTP mix | 2.25 |
| MGSO4 (50mM) | 1.5 |
| Primer (50µM) - crt F | 0.5 |
| Primer (50µM) - crt R | 0.5 |
| Primer (50µM) - dhfr F | 0.6 |
| Primer (50µM) - dhfr R | 0.6 |
| Primer (50µM) - dhps F | 1 |
| Primer (50µM) - dhps R | 1 |
| Primer (50µM) - mdr1 F | 0.6 |
| Primer (50µM) - mdr1 R | 0.6 |
| Primer (50µM) - k13 F | 0.8 |
| Primer (50µM) - k13 R | 0.8 |
| Primer (50µM) - csp F | 1 |
| Primer (50µM) - csp R | 1 |
| DNA Pol | 0.5 |
| gDNA | 4 |

| | |
|---|---|
| H2O | 26.75 |
| Total | 50 |

All of the samples described in this study underwent multiplex drug resistance and *csp* amplification using the Thermo Fisher Platinum™ *Pfx* DNA Polymerase (11708039). Note that the quantity of primers added for each reaction varied according to the reaction efficiency, to attempt to make amplicon coverage more even. The Platinum™ *Pfx* DNA Polymerase enzyme has been discontinued by the manufacturer. We have found that the Kapa HiFi polymerase (KK2101, manufacturer conditions) produces comparable results using the same primers.

The reaction conditions for the drug resistance and *csp* multiplex PCR using *Pfx* DNA Polymerase, used for leucodepeleted venous blood samples, are shown below:

| **Step no.** | **Step** | **Temp** | **Duration** |
|---|---|---|---|
| 1 | Initial denature | 94 | 5 min |
| 2 | Denature | 94 | 15 sec |
| 3 | Anneal | 54 | 30 sec |
| 4 | Extend | 68 | 1 min |
| 5 | Repeat steps 2-4 34x more times | | |
| 6 | Store | 4 | Forever |

The drug resistance and *csp* multiplex PCR with *Pfx* DNA Polymerase used for dried blood spot (DBS) samples was the same as that used for leucodepleted venous blood samples (above) except for the following modifications: 15ul of extracted gDNA was used as template for the PCR reaction (with accordingly reduced H2O added to reach 50ul reaction total), and 40 PCR cycles was used instead of 35 cycles (i.e. step 5 was repeated 39 more times instead of 34).

The reaction mix for the *msp1* PCR is shown below:

| | **1x** |
|---|---|
| GoTaq MM | 25 |
| msp1 F (10uM) | 1 |
| msp1 R (10uM) | 1 |
| gDNA | 2 |
| H2O | 21 |
| Total | 50 |

The reaction conditions for the *msp1* PCR are shown below:

| **Step no.** | **Step** | **Temp** | **Duration** |
|---|---|---|---|
| 1 | Initial denature | 95 | 2 min |
| 2 | Denature | 93 | 20 sec |
| 3 | Anneal | 54 | 25 sec |
| 4 | Extend | 65 | 5 min |
| 5 | Repeat steps 2-4 34x more times | | |

| | | | |
|---|---|---|---|
| 6 | Final extend | 72 | 10 min |
| 7 | Store | 4 | Forever |

The *msp1* PCR was performed using Promega long-range GoTaq® Polymerase (M4021).

#### qPCR to estimate human and *P. falciparum* gDNA relative abundance

Quantitative PCR (qPCR) was used to assess the ratio of *P. falciparum* to human DNA present in the mock clinical DBS samples and the clinical DBS samples collected in Navrongo. Following DNA extraction, qPCR was performed on a Roche LightCycler®480 Instrument II with a fluorescent probe method using distinct dyes FAM, HEX and RED 640. The primer/probes from Integrated DNA Technologies (IDT) were designed on PrimerBLAST; AMA1 for *P. falciparum*, PVX_096055 for *Plasmodium vivax* (VIV3) and PDGFRB for *Homo sapiens* (PLAT1). Probe dyes; FAM for VIV3, LightCycler 640 for AMA1 and HEX for PLAT1. The addition of an internal quencher, ZEN was used for both HEX and FAM dyes. The *P. vivax* probe played no role in this study. Primer sequences are shown below.

| Name | Type | Primer sequence |
|---|---|---|
| VIV3 | Primer-forward | AAA GAT TCG TAG CTG TCG GTG GGT |
| | Primer-reverse | TTC CAT TAA GTG CGC GTA CCG AGA |
| | Probe | ACA GCG ACG ACT CCA GAT CCG ATT TA |
| AMA1 | Primer-forward | TGC CAT ATA TTC CGT CCA TGG |
| | Primer-reverse | ACG CAT ATC CAA TAG ACC ACG |
| | Probe | CGA ACC CGC ACC ACA AGA ACA AAA |
| PLAT1 | Primer-forward | CTT ACC ACA TCC GCT CCA TC |
| | Primer-reverse | TTC ACA CTC TCC GTC ACA TTG |
| | Probe | CAC ATC CCC AGT GCC GAG TTA GA |

A master mix was prepared with the following components (quantities provided per well); 10µl Roche LightCycler®480 Probes Master (04887301001), 6µl Nuclease-free water (AM9937) 1µl AMA1 Primer/probe, 1µl PLAT1 Primer/probe, and 1µl VIV3 Primer/probe. Sample volumes: 1µl sample DNA, 1 µl negative control and 1 µl positive controls (standards), with all samples including a repeat.

Program selection was 'Abs quant'. PCR conditions are shown below:

| Step | Temperature | Duration | Cycles |
|---|---|---|---|
| 1 | 94°C | 5 minutes | |
| 2 | 57°C | 20 seconds | Steps 2-4: Total of 45 cycles |
| 3 | 72°C | 1 second | |
| 4 | 95°C | 10 seconds | |
| 5 | 40°C | infinite | |

qPCR results for the mock DBS samples are shown in **Supplementary Figure 1**, and for the clinical DBS samples collected in Navrongo in Supplementary Figure 6.

#### Inference rules for drug resistance phenotyping from genotype data

Inference rules were based on Jacob, C., *et al*. 2021. *eLife* 10:e62997, summarised below:

| **Antimalarial drug** | **Gene** | **Mutation** | **Interpretation** |
|---|---|---|---|
| Chloroquine (CQ) | *crt* | 76-K<br>76-T | Susceptible<br>Resistant |
| Pyrimethamine (PYR) | *dhfr* | 108-S<br>108-N | Susceptible<br>Resistant |
| Sulfadoxine (SX) | *dhps* | 437-A<br>437-G | Susceptible<br>Resistant |
| Sulfadoxine-Pyrimethamine (SP) for malaria treatment | *dhfr* | 51-N or 59-C or 108-S<br>51-I & 59-R & 108-N | Susceptible<br>Resistant |
| Sulfadoxine-Pyrimethamine (SP) for IPTp | *dhfr* & *dhps* | *dhfr*: 51-N or 59-C or 108-S<br>& (*dhps*: 437-A or 540-K)<br>& (*dhfr*-164-I or *dhps*-581-A or *dhps*-613-A)<br><br>*dhfr*: 51-N & 59-C & 108-S<br>& *dhps*: 437-G & 540-E<br>& (*dhfr*-164-L or *dhps*-581-G or *dhps*-613-(S/T)) | Susceptible<br><br><br><br>Resistant |
| Artemisinin | *kelch13* | Mutations associated with artemisinin partial resistance* | Artemisinin partial resistance |

*Crt = chloroquine resistance transporter,* PF3D7_0709000; *dhfr = dihydrofolate reductase-thymidylate synthase,* PF3D7_0417200; *dhps = dihydropteroate synthetase,* PF3D7_0810800; *kelch13* = PF3D7_1343700. IPTp = Intermittent preventive therapy in pregnancy. *The *kelch13* mutations that may be associated with artemisinin partial resistance were taken from the MalariaGEN technical documents, accessed 2022-08-07, and are listed below:

- P441L, F446I, G449A, D452E, N458Y, C469Y, C469F, M476I, K479I, A481V, Y493H, R515K, S522C, P527L, N537I, N537D, G538V, R539T, I543T, P553L, R561H, V568G, P574L, R575K, M579I, C580Y, D584V, P667T, F673I, A675V, H719N
